# Two CENH3 paralogs in the green alga *Chlamydomonas reinhardtii* have a redundantly essential function and associate with ZeppL-LINE1 elements

**DOI:** 10.1101/2025.03.28.645958

**Authors:** Dianyi Liu, Mingyu Wang, Jonathan I. Gent, Peipei Sun, R. Kelly Dawe, James Umen

## Abstract

Centromeres in eukaryotes are defined by the presence of histone H3 variant CENP-A/CENH3. Chlamydomonas encodes two predicted CENH3 paralogs, CENH3.1 and CENH3.2, that have not been previously characterized. We generated peptide antibodies to unique N-terminal epitopes for each of the two predicted Chlamydomonas CENH3 paralogs as well as an antibody against a shared CENH3 epitope. All three CENH3 antibodies recognized proteins of the expected size on immunoblots and had punctuated nuclear immunofluorescence staining patterns. These results are consistent with both paralogs being expressed and localized to centromeres. CRISPR-Cas9 mediated insertional mutagenesis was used to generate predicted null mutations in either *CENH3.1* or *CENH3.2*. Single mutants were viable but *cenh3.1 cenh3.2* double mutants were not recovered, confirming that the function of CENH3 is essential. We sequenced and assembled two chromosome-scale Chlamydomonas genomes from strains CC-400 and UL-1690 (a derivative of CC-1690) with complete centromere sequences for 17/17 and 14/17 chromosomes respectively, enabling us to compare centromere evolution across four isolates with near complete assemblies. These data revealed significant changes across isolates between homologous centromeres including mobility and degeneration of ZeppL-LINE1 (ZeppL) transposons that comprise the major centromere repeat sequence in Chlamydomonas. We used Cleavage Under Targets and Tagmentation (CUT&Tag) to purify and map CENH3-bound genomic sequences and found enrichment of CENH3-binding almost exclusively at predicted centromere regions. An interesting exception was chromosome 2 in UL-1690, which had enrichment at its genetically mapped centromere repeat region as well as a second, distal location, centered around a single recently acquired ZeppL insertion. The CENH3-bound regions of the 17 Chlamydomonas centromeres ranged from 63.5kb (average lower estimate) to 175kb (average upper estimate). The relatively small size of its centromeres suggest that Chlamydomonas may be a useful organism for testing and deploying artificial chromosome technologies.

**Significance statement:** Histone H3 variant CENP-A/CENH3 is a conserved centromere protein in eukaryotes but has not been well-characterized in green algae. Our data establish the function of CENH3 at Chlamydomonas centromeres and identify an unexpected strain-specific neocentromere candidate region in chromosome 2 likely caused by a single ZeppL-LINE1 insertion into a genic region that is distal to the native chromosome 2 centromere.

## Introduction

Centromeres are defining features of chromosomes in most eukaryotes and are essential for ensuring precise mitotic and meiotic genome segregation. In many species including plants and animals, the centromeric region of each chromosome is specified epigenetically and is marked by a specialized chromatin environment. These chromatin specializations allow assembly and maintenance of kinetochores that mediate mitotic or meiotic spindle microtubule attachment and chromosome segregation (Sullivan and Karpen 2004). The DNA in centromeric regions is often characterized by large arrays of transposon or satellite DNA repeats, but such repeats are insufficient to specify a new centromere (Hartley and O’Neill 2019; Miga 2019). Instead, specific centromeric proteins are present at centromere regions and can recruit new centromeric proteins to maintain centromere identity (Earnshaw 2015; Yoda et al. 2000).

A key protein of centromeres is the histone H3 variant CENP-A/CENH3 which incorporates into centromeric nucleosomes in place of canonical histone H3 (Sundararajan and Straight 2022). CENH3 not only marks the location of the centromere but serves as the base of the larger kinetochore complex (Kixmoeller, Allu, and Black 2020). Overexpression or ectopic localization of CENH3 can be sufficient to drive formation of a new centromere (Furuyama and Biggins 2007; Feng et al. 2020). Not surprisingly, *CENH3* has proven to be an essential gene in all species where it has been carefully analyzed H3s (Earnshaw 2015; Sundararajan and Straight 2022).

CENH3 homologs have been identified in many species and have some unusual properties compared with canonical histone. These properties include a highly variable N-terminal tail region and substitutions at a few positions within the highly conserved histone fold domain. Unlike canonical histone H3 which shows little variability across large phylogenetic distances, there is limited sequence conservation of CENH3 specific sequences between species outside the conserved histone fold domain, with evidence of positive selection on CENH3 within some clades (Draizen et al. 2016). The rapid evolution of CENH3 proteins is mirrored by high rates of change between species in centromeric repeat sequences (Talbert and Henikoff 2022).

At ∼113 Mb, the *Chlamydomonas reinhardtii* (Chlamydomonas) genome is one of the smallest among photosynthetic eukaryotes (Craig et al. 2021; Payne et al. 2023). The Chlamydomonas nuclear genome has 17 chromosomes with the smallest being only ∼3.8 Mb (Craig et al. 2021), closer in size to Chromosome IV of yeast (∼1.5 Mb) than the smallest Arabidopsis chromosome (∼26 Mb) (Hou et al. 2022; Naish et al. 2021). Like yeast, Chlamydomonas can be propagated like a microbe, subjected to rapid genetic manipulation, and grown at industrial scale (Scaife et al. 2015). Chlamydomonas is an important model species for research in photosynthesis, cell cycle control, metabolism, and biotechnology applications (Strenkert et al. 2019; Salomé and Merchant 2019). The utility of Chlamydomonas in both basic and applied research combined with its relatively small genome size and compact individual chromosomes makes it a particularly suitable model for chromosome-scale synthetic biology applications.

Any effort to carry out whole chromosome design in Chlamydomonas will require a more complete understanding of the centromeres. Centromere locations in Chlamydomonas have been estimated genetically (Kathir et al. 2003; Lin, Cliften, and Dutcher 2018) and subsequently sequenced, revealing the presence of nested Zepp-like transposons, a class of L1 LINE elements found in Chlamydomonas relatives and other green algae (Craig et al. 2021). Based on the lengths of ZeppL-LINE1 (ZeppL) clusters, the centromere sizes in a complete genome of the CC-5816 strain were estimated to range from 252 to 480 kb (Payne et al. 2023). However, centromeres in most species are defined not by DNA sequence, but by the presence of CENH3, which may occupy only a subset of the repeat region at each centromere (Sundararajan and Straight 2022). In Arabidopsis, the major centromere repeat *AthCEN178* occupies regions extending over 1.5-6.5 Mb, but the CENH3-interacting regions are only ∼1-2 Mb (Wlodzimierz et al. 2023). These and similar data from other plant species suggest that the functional centromere regions of Chlamydomonas are likely to occupy only a fraction of the sequence within the ZeppL clusters.

Here we describe the two *CENH3* paralogs in Chlamydomonas, *CENH3.1* and *CENH3.2*, and demonstrate that predicted null mutants for either gene alone are viable but that a *cenh3.1 cenh3.2* double mutant is inviable. We generated chromosome-scale assemblies of Chlamydomonas strains UL-1690 and CC-400 and compared homologous centromere regions among four strains with largely contiguous assemblies across their centromeres, and uncovered evidence of recent ZeppL element mobility and degeneration. We next identified CENH3.1 and CENH3.2 bound genomic regions in strain UL-1690 using CUT&Tag which validated the predicted centromere localization of the two proteins and allowed us to annotate functional centromere domains. The data show that centromeres in Chlamydomonas are relatively small, occupying on average ∼140 kb regions embedded within centromeric ZeppL clusters (Craig et al. 2021, 2023). Unexpectedly, a small non-centromeric locus on chromosome 2 of strain UL-1690 contained peaks of both CENH3.1 and CENH3.2 binding centered around a single strain-specific ZeppL insertion, hinting at a potential mechanism for new centromere formation.

## Results

### Validation of Chlamydomonas CENH3 paralogs

Chlamydomonas is predicted to encode two *CENH3* paralogs, *CENH3.1* (Cre16.g661450) and *CENH3.2* (Cre02.g104800) respectively, with the *CENH3.2* gene model correctly predicted in the Phytozome V6.1 assembly and annotations (Goodstein et al. 2012), but not in V5.6 which mis-predicted its structure. These paralogs were identified based on their divergence from canonical histone H3 sequences in specific N- and C-terminal regions, and other features that are common to CENH3 proteins (Ravi et al. 2010). Previous transcriptome analyses in synchronous wild-type cultures showed that both genes were expressed periodically with peaks during S/M phase along with other replication-dependent histone genes (Figure S1a). However, the transcript abundance of both *CENH3* paralogs was much lower than those for canonical histone genes, and the peak transcript abundance of *CENH3.1* was around three-fold higher than that of *CENH3.2* (Figure S1b) (Zones et al. 2015; Strenkert et al. 2019).

Three Chlamydomonas-specific CENH3 specific antibodies were raised against peptides that contained unique paralog-specific sequences for CENH3.1 and CENH3.2, and a peptide covering a region that is conserved between CENH3.1 and CENH3.2, but not found in other histone H3 sub-types (Figure 1a, Materials and Methods). All three custom CENH3 antibodies recognized proteins of the expected size (∼ 17kD) on immunoblots, as well as some nonspecific cross-reacting proteins (Figures 1b, S1c). All three antibodies had similar punctate nuclear immunofluorescence (IF) staining patterns as expected for centromere localized antigens (Figures 1c, S1d). While we cannot completely rule out cross reactive antigens as the source of the IF staining patterns, the similar nuclear staining patterns for all three antibodies and additional evidence showing CENH3.1 and CENH3.2 at centromeric regions (see below CUT&Tag results) suggests the puncta represent centromeres.

**Figure 1.**
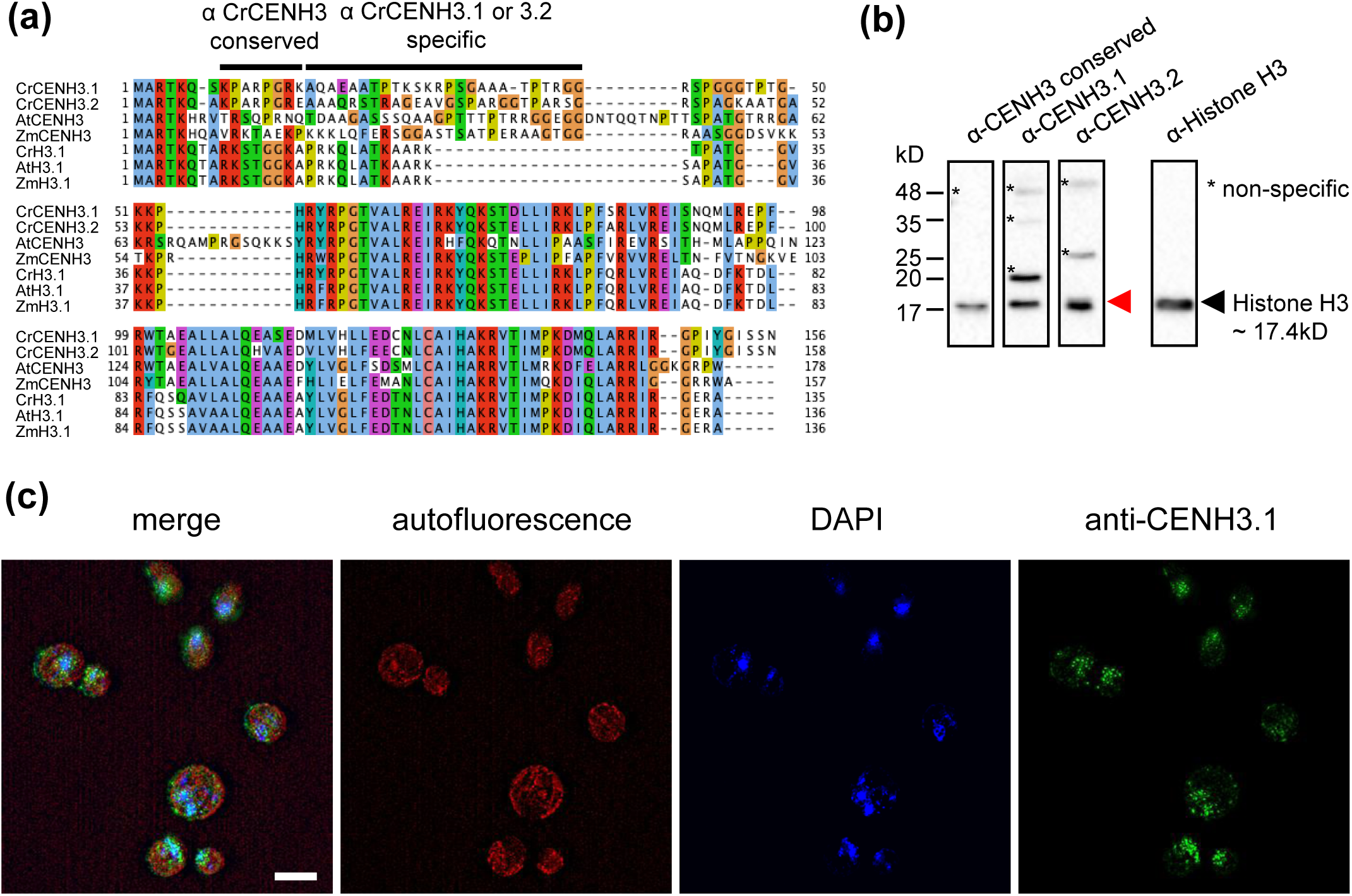
Identification of two Chlamydomonas CENH3 paralogs. **(a)** Multiple sequence alignment of CENH3 and conventional histone H3 proteins from Chlamydomonas (Cr), *Arabidopsis thaliana* (At) and *Zea mays* (Zm) (Khan *et al*., 2018; Cui *et al*., 2015). The alignment is colored to show conserved residues. Lines above the alignments show peptide sequences from CrCENH3.1 or CrCENH3.2 to which antibodies were raised and purified. **(b)** Immunoblots of SDS-PAGE separated protein lysates of wild type using α-CENH3.1-specific, α-CENH3.2-specific, α-CENH3-conserved, or α-histone H3 (internal loading control). The black arrowhead shows the position of histone H3. The red arrowheads show the position of CENH3 proteins. Asterisks mark cross-reacting non-specific antigens. **(c)** Immunofluorescence microscopy images of wild-type UL-1690 immunostained with CENH3.1-specific antibodies (pseudo colored magenta) and stained with DAPI (pseudo colored cyan). Background autofluorescence and residual chlorophyll is pseudo colored red. Merge contains an overlay of the three channels. The lower row is a higher magnification inset from the indicated region. Scale bar = 5 μm.

### CENH3.1 and CENH3.2 are essential but functionally redundant

We used CRISPR-Cas9 mediated insertional mutagenesis (Shin et al. 2016; Picariello et al. 2020) to generate predicted null alleles for each of the two Chlamydomonas *CENH3* paralogs in a wild-type strain background, UL-1690 (Figure 2a) (Materials and Methods).

**Figure 2.**
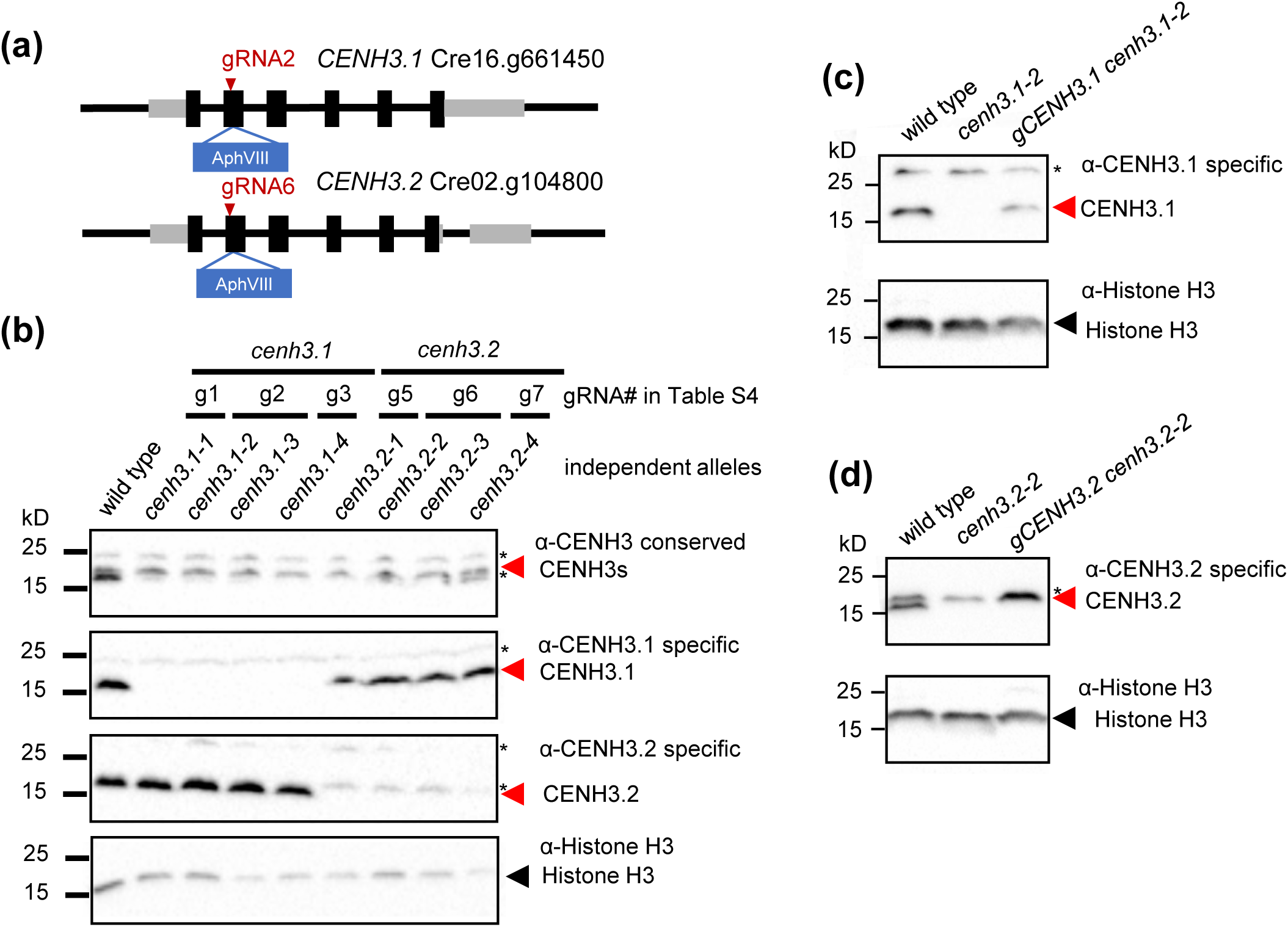
CrCENH3.1 and CrCENH3.2 are functionally redundant. **(a)** Schematic of *CENH3.1* and *CENH3.2* loci and Phytozome v6.1 IDs with location of targeted *AphVIII* marker insertion guided by CRISPR-Cas9 cutting at the indicated gRNA sites. Black rectangles, exons; dark green rectangles, untranslated regions; black lines, introns, and intergenic regions. Red arrows show target sites of gRNA used to generate *cenh3.1-2* and *cenh3.2-2* respectively. Inserted *AphVIII* selectable marker conferring paromomycin resistance is in blue. Note that the *CENH3.1* (Cre16.g661450) gene model was missing UTRs in Phytozome V6.1, so the UTR coordinates were taken from V5.6 gene predictions. The *CENH3.2* (Cre02.g104800) gene model was correctly predicted in V6.1, but not in V5.6. **(b)** Immunoblots of SDS-PAGE separated protein lysates of wild type or independent *cenh3* mutants (g1, g2, g3 and g5, g6, g7) detected using α-CENH3.1-specific, α-CENH3.2-specific, α-CENH3-conserved, or α-histone H3 (internal loading control) antibodies. The black arrowhead shows the position of histone H3. The red arrowheads show the position of CENH3 proteins. Asterisks mark cross-reacting non-specific antigens. Note that the α-CENH3-conserved immunoblot signal is reduced and somewhat weak in the mutant strains **(c,d)** Immunoblots of transgenic rescue strains for **(c)** *cenh3.1-2* or **(d)** *cenh3.2-2*. Blots are labeled as in panel **(b)**.

Multiple insertional alleles were generated and confirmed for each paralog indicating that neither of the two genes individually have essential functions (Materials and Methods, Table S1-S2).

Immunoblotting with each of the paralog-specific antibodies was used to validate loss of the predicted CENH3 protein in mutant strains (Figure 2b). Two null strains containing marker insertions in their second exons were used for all further experiments and referred to as *cenh3.1-2* and *cenh3.2-2* respectively (Figures 2a, S2a, Table S1). Immunoblots using the antibody raised against the common epitope of CENH3.1 or CENH3.2 had reduced signal in single mutant strains suggesting a lack of compensatory upregulation of the intact paralog when the other paralog was missing (Figure 2b). Neither of the *cenh3* mutants had obvious growth or viability phenotypes, and both showed normal Mendelian segregation in progeny from backcrosses to a wild-type strain (Figures S2b, S2c). However, when a *cenh3.1-2* mutant was crossed with a *cenh3.2-2* mutant, *cenh3.1-2 cen3.2-2* double mutants were not recovered among the progeny, indicating that the two paralogs redundantly carry out an essential function (Figure S2d).

Constructs with native genomic *CENH3.1* or *CENH3.2* genes (Figure S2e) were introduced into corresponding *cenh3.1* or *cenh3.2* mutant strains, respectively, and screened for restoration of expression (Materials and Methods). Candidate rescued strains were first screened by genotyping, then followed by direct screening for restoration of CENH3 signal on immunoblots (Materials and Methods, Figures 2c, 2d). Confirmation of transgene function was obtained by crossing each rescued strain to a mutant strain with a *cenh3* allele in the non-rescued paralog. Progeny from the cross were screened for each of the two mutant *cenh3* alleles and for the presence of the rescuing transgene. Unlike the intercross of *cenh3* mutants without a rescue construct, *cenh3.1-2 cenh3.2-2* double mutant progeny could be recovered when a rescuing *CENH3.1* or *CENH3.2* transgene was also present (Figure S2f), indicating that the transgenes expressed functional CENH3.

### High-quality genome and centromere assemblies of UL-1690 and CC-400

Laboratory strains UL-1690 (a derivative of strain CC-1690/21gr) and CC-400 (cw15) were selected for HiFi PacBio sequencing to obtain contiguous assemblies of centromere regions (Materials and Methods). CC-1690 is a standard wild-type laboratory strain (Sager 1955; Pröschold, Harris, and Coleman 2005), but sequence comparisons (see below) suggested that our isolate was likely the progeny of an intercross between CC-1690 and another strain. To avoid confusion, we designated our CC-1690 isolate as UL-1690 (Umen Laboratory 1690). CC-400 is a commonly used wall-deficient mutant strain background that has been separated from a common ancestor with CC-1690 for around 80 years (Pröschold, Harris, and Coleman 2005; Hyams and Davies 1972; Roy Davies and Plaskitt 1971). The original CC-1690 genome was assembled previously, but with a high base-error rate due to limitations of Nanopore sequencing (O’Donnell et al., 2020). The genomes of our strains UL-1690 and CC-400 were assembled using PacBio HiFi long read sequencing data with hifiasm as the assembler (Cheng et al. 2021) and RagTag for scaffolding (Alonge et al. 2022) (Materials and Methods). We assessed the accuracy of the assemblies by comparing them to a recent gapless assembly of the strain CC-5816 (Figure S3a, S3b) (Payne et al. 2023). In the UL-1690 assembly, six chromosomes are gapless, with 25 gaps across the remaining 11 chromosomes. The CC-400 assembly also has six gapless chromosomes and 16 gaps across the other 11 chromosomes (details see Table S3). The CC-4532 strain serves as the standard *Chlamydomonas reinhardtii* reference genome with ongoing updates (Craig et al. 2023). In the current CC-4532 assembly (Phytozome V6.1), 63 gaps remain, and only six of 17 centromeres (chr4, 5, 7, 9, 14, 16) are fully assembled. In contrast, our new HiFi assemblies include 14 fully assembled centromeres in UL-1690 (chr1, 2, 4-7, 9-11, 13-17), and all 17 centromeres in CC-400 (Table S3).

The CC-4532 and CC-1690 genome assemblies differ from UL-1690, CC-400 (this manuscript) and CC-5816 (Payne et al) by the presence of several large tandem repeat blocks, most noticeable on chr11 and chr15 in the latter three assemblies but not the former two (Figure S4). The chr11 block is a single ∼670 kb (UL-1690) or ∼860 kb (CC-5816, CC-400) insertion comprising over three thousand copies of a 184 bp repeat, while the chr15 repeat insertions are in multiple locations and involve several repeat sequences with repeat units ranging from 107 to 438 bp and occupying a total of around 1.2 Mb on chr15 in all three strains (Figure S4, Table S4). Like centromere regions, these repeats are nearly devoid of predicted protein coding genes, highly enriched for 5-methylcytosine (Payne et al. 2023) and may form heterochromatin-like domains, but this remains to be determined.

### Functional centromere size estimates based on CUT&Tag analysis of CENH3

Centromeres in most eukaryotes are defined by the presence of CENH3 which is maintained at centromere repeats through an epigenetic recruitment mechanism (Feng et al. 2020). Centromere location and size are best estimated by determining the sub-regions of centromere repeats bound to CENH3. To identify these regions in Chlamydomonas, we carried out Cleavage Under Targets and Tagmentation (CUT&Tag) (Kaya-Okur et al. 2019, 2020) with our sequenced strain UL-1690 using each of the three CENH3 antibodies described above, along with non-specific rabbit IgG as a control for background signal. The CENH3 CUT&Tag data from each of the three antibodies showed nearly identical peak locations, with localization confined almost exclusively to ZeppL elements at centromere regions for all 17 Chlamydomonas chromosomes (Figure 3, Materials and Methods).

**Figure 3.**
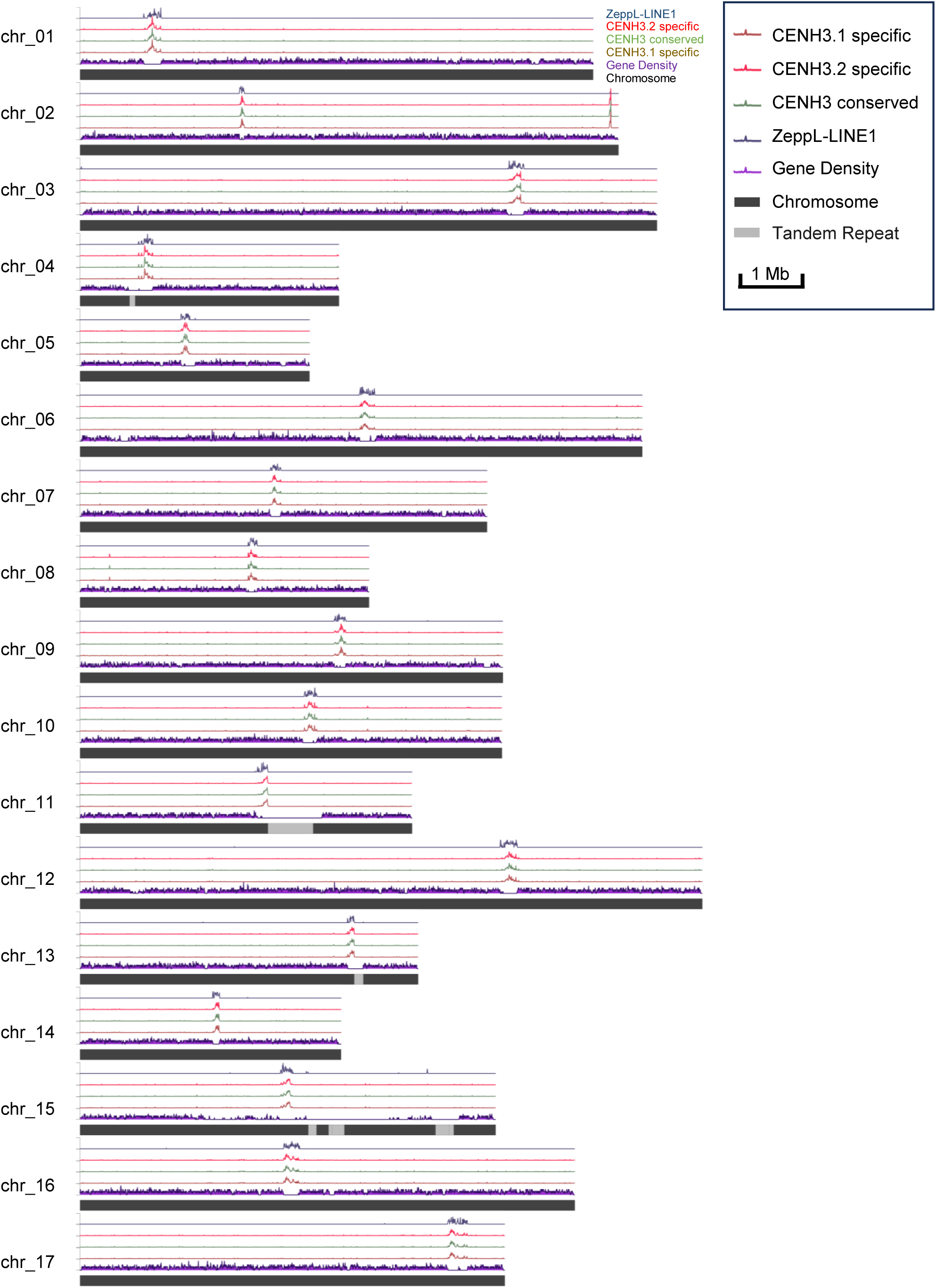
CENH3 localization in the UL-1690 genome assembly. For each chromosome (represented as a black bar), six tracks from bottom to top indicate protein coding gene density, ZeppL repeats, and normalized CUT&Tag enrichment (multiple plus unique mapping) for CENH3.1-specific, CENH3.2-specific and CENH3-conserved antibodies. Gray regions of the assembly represent large repeat blocks missing from the CC-4532 V6.1 reference assembly hosted on Phytozome. The y axes for fold enrichment of CENH3.1-specific, CENH3.2-specific, and CENH3-conserved peaks range from 0 to 75. The y axes for ZeppL-LINE1 enrichment and gene density plot ranges from 0 to 15 and 0 to 6. The window size for all tracks is 5kb.

Prior data from multiple species showed that distribution of CENH3 reads is usually denser in the center of centromeres and decreases toward chromosome arms, and that the pattern of CENH3 enrichment shifts between individuals (Gent, Wang, and Dawe 2017). Thus, our data represents a population average. Additionally, the highly repetitive nature of centromeric DNA often makes it impossible to know the precise origin of a read. Alignment parameters that allow reads to map to one of multiple possible locations (as well as unique locations, MAPQ value of 20 or larger) tend to overestimate the centromere DNA occupied by CENH3, likely beyond the minimal regions required for centromere function. In contrast, using only unique mapping reads provides more accurate estimates of the locations of functional centromeres but may underestimate their total size. Here, we utilized both multiple plus unique mapping and unique mapping to estimate the upper and lower centromere size limits for each chromosome (Figure 4a, Table 1). The results showed that the upper estimates of centromere sizes are 65 to 310 kb, which is close to the size of ZeppL enriched regions (Table 1, Figure S5a, Table S5), and the lower estimates of centromere sizes are 50 to 105 kb with ∼85% of unique mapping regions containing ZeppL sequences (Table 1, Figure S5b, Table S5).

**Figure 4.**
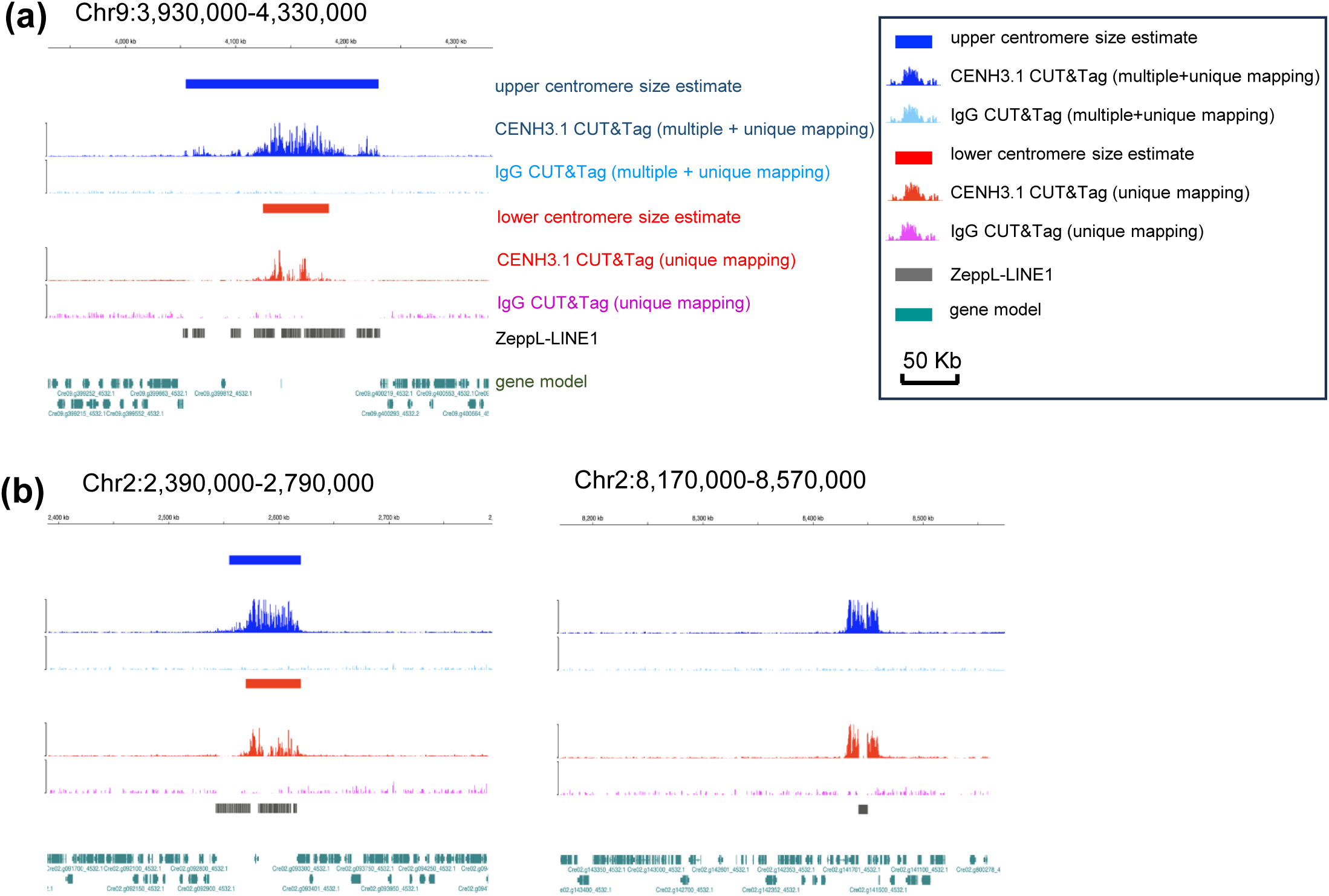
Genome browser views of selected centromere regions showing details of CENH3.1 CUT&Tag read distribution. Blue or red bars above CENH3 peaks show upper (blue) and lower (red) size estimates for the centromere based on combined multimapping plus unique mapping reads or only unique mapping reads, respectively (see Materials and Methods for details). **(a)** Example of an averaged-size centromere from chr9. **(b)** Left panel, genetically defined centromere region for chr2. Right panel, possible neocentromere region on chr2 (see main text for details).

**Table 1.** Chromosome length, ZeppL-LINE1 region length, and centromere size estimates for UL-1690 HiFi genome assembly.

Chromosome sizes and ZeppL-enriched region sizes exhibited a weak positive correlation (R^2^ = 0.21, Figure S5c), but when uniquely mapping centromere regions from our lower centromere size estimates were compared to chromosome sizes the correlation largely disappeared, suggesting that functional centromere size may be independent of chromosome size in Chlamydomonas.

### Possible secondary centromere on Chromosome 2

Chlamydomonas has monocentric chromosomes, where the functional centromeres are limited to a single domain containing CENH3. However, on chromosome 2 we observed a second strong peak of CENH3 CUT&Tag read enrichment far from its mapped centromere and close to the end of its long arm (Figures 3, 4b), suggesting a possible secondary centromere.

The total estimated size of the secondary CENH3 binding region is 30-35 kb, which is about half the size of the genetically defined centromere on chromosome 2 (Table 1). Remarkably, a single intact ZeppL element (described above) lies at the center of the CENH3 enriched region, in between two protein coding genes Cre02.g800275_4532.1 and Cre02.g141600_4532.1 (Figure S6a). Importantly, the CENH3 CUT&Tag signal is strongly enriched not only in the ZeppL sequences, but in the unique gene-rich sequences flanking the ZeppL insertion, indicating the unambiguous presence of CENH3 at this locus (Figures 4b, S5a). The ZeppL insertion near the end of chromosome 2 was unique for strain UL-1690 and not found in assemblies of CC-4532, CC-5816, CC-400, or the previously assembled Nanopore CC-1690 genome (O’Donnell et al., 2020) (Figure S6b, Table S6). These results suggest that the ZeppL element near the end of chromosome 2 in UL-1690 could be the result of a recent transposition event.

### Evidence of active ZeppL transposition and turnover in Chlamydomonas

The sizes of the predicted centromere regions for a given chromosome (measured as distance between the ZeppL repeats flanking the centromere region) are similar between strains, suggesting that overall activity of ZeppL is not high enough to cause major changes to centromere size over the decades separating CC-400, CC-1690 and CC-5816 (Pröschold, Harris, and Coleman 2005) (Table S6, Data S1). However, the finding of the likely ZeppL transposition activity associated with insertion at the neocentromere-like region on Chromosome 2 prompted us to search for the origin of the insertion element and for additional signs of ZeppL mobility. We first identified 49 full length or near full length ZeppL sequences (>8 kb) as potential active candidates among four genome assemblies (UL-1690, CC-400, CC-1690, CC-5816) and eleven different chromosomes (1,2,3,6,9,10,13,14,15,16,17) including the insertion on Chr2 (Data S2). We also found four partial ZeppL sequences in CC-400 and UL-1690 centromeres which were >5kb and >99.9% identical to the corresponding full length ZeppL element in strain CC-5816. These four elements were likely beginning a process of fragmentation and decay (Data S3, Table S7). The phylogeny of the full-length sequences revealed two separate sub-groups that we categorized as relatively stable ZeppL elements and recently active ZeppL elements (Figures 5, S7-S9). The relatively stable ZeppL elements formed distinct and well-supported chromosome-specific clades in the centromeres of chr1 (two elements), 6, 9, 10, 13, 14, 15, 16. These elements were present in all four strains though there was evidence of degeneration/fragmentation of the originally full-length ZeppL element as described above (Figure 5, Data S3, Table S7). The recently active sub-group of ZeppL elements were present on chromosomes 1, 2, 3 (two elements),10,15,16,17 and included the neocentromere candidate ZeppL sequence on chromosome 2 of UL-1690. This group was characterized by their high relatedness (near-zero branch lengths) suggesting more recent origins via active transposition between chromosomes. The branching patterns of the recently active ZeppL group to which the new insertion on chromosome 2 belongs suggests some of these elements were present in the recent common ancestor of all four laboratory strains for which we have data. However, there also appears to have been active movement evidenced by presence/absence between strains for elements on chromosomes 1, 10 and 17 (Figure 5). In summary, a group of highly similar ZeppL elements shared across multiple chromosomes, including the neocentromere of chr2, suggests active ZeppL transposition and some element decay before and since the four laboratory strains separated.

**Figure 5.**
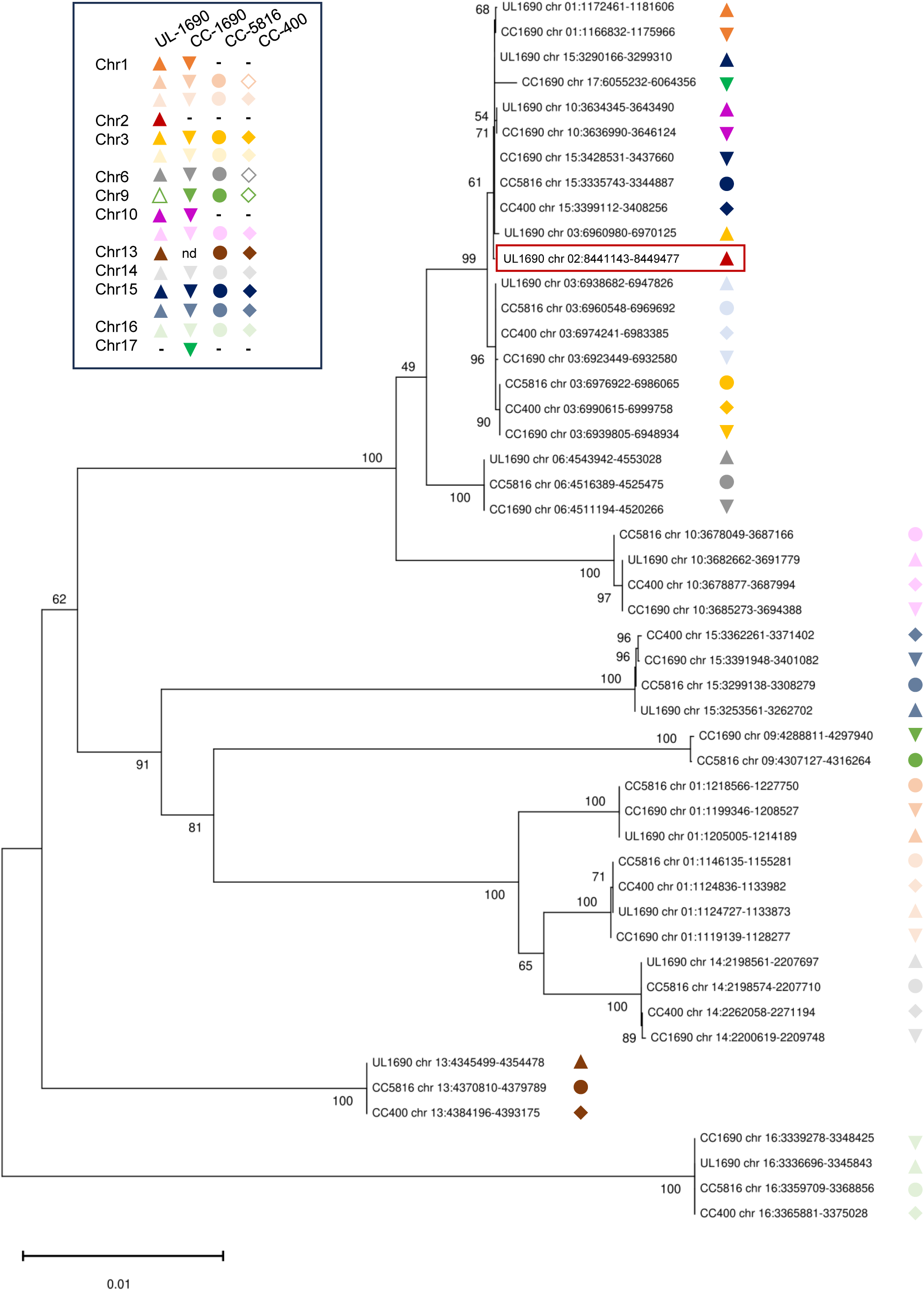
Phylogenetic analysis of near full length ZeppL sequences (>8kb) in four genome assemblies. Unrooted Neighbor Joining tree of 49 full length or near full length ZeppL elements in four strains. Bootstrap values are shown at each node. Each shape (square, circle, triangle, diamond) represents a specific strain. Each color represents a specific ZeppL element. Filled shapes indicate an intact ZeppL element and hollow shapes represent a partially decayed copy (between 5kb and 8kb) corresponding to a full length ZeppL element at the same location in other strains. Dashes indicate absence of a ZeppL element at that location in one or more strains. nd indicates no data due to an assembly gap in that strain. The lower portion of the tree contains distinct, centromere-specific ZeppL elements found in the same position within the centromere in all strains where there was a gapless assembly at that centromere. The upper part of the tree contains a group of highly similar, recently active ZeppL elements (near zero branch lengths) shared across several chromosomes, including the newly inserted element in chr2 (boxed; see main text for details). Full length ZeppL sequences and coordinates can be found in Data S2. Partly decayed ZeppL sequences and coordinates can be found in Data S3.

### Centromere variability between and within Chlamydomonas strains

The identification of ZeppL element mobility led us to compare centromere lengths and sequences more broadly among different strains and assemblies. Overall individual centromere repeat region lengths and sequences were similar between different laboratory strains (Figures S10-S12), similar to what has been observed in other species (Langley et al. 2019; Altemose et al. 2022; Liu and Dawe 2023; Wlodzimierz et al. 2023). However, there were also some notable differences caused by structural rearrangements (Figure S13). While it is difficult to meaningfully compare different centromeres with conventional sequence alignments due to their heterogeneity in size and sequence, alignment-free methods based on k-mer counting can be used to compare centromere relatedness across all centromeres and strains (Zielezinski et al. 2017; Luczak, James, and Girgis 2019).

We used Mash (Ondov et al. 2016) to estimate Jaccard distances between each of 64 gap-free centromere sequences available from four genome assemblies (UL-1690, CC-1690, CC-400 and CC-5816) (Materials and Methods) and visualized their divergence estimates with a distance tree (Figure 6). As expected, homologous centromeres formed distinct clades and showed varying degrees of relatedness with each other. The chr4 and chr10 centromeres were most closely related to each other while the chr2 native centromere was on average most distal to all other centromeres. For each specific centromere the amount of inter-strain divergence also varied by chromosome. Most individual centromeres for a given chromosome were highly similar in all four strains indicated by short branch lengths connecting them to their nearest common node; but in a few cases there was marked divergence in one or more of the strains e.g. CC-1690 chr1; CC-400 chr10; CC-400, CC-1690 chr14; CC-400 chr15. For chr10 and chr15 one centromere border differed between CC-400 and the other three strains, including a ∼200 kb inversion in CC-400 that substantially changed the border of its chr15 centromere (Figure S13). For chr1 and chr14 there were internal strain-specific indels in CC-1690 and or CC-400 that were likely responsible for higher divergence scores of their centromeres from the other strains (Figures S7-S9). In summary, comparative analyses indicate that both large-scale genomic changes—such as structural rearrangements and ongoing transposon activity—and smaller-scale polymorphisms have driven the rapid and continuous evolution of centromeres, even within the relatively short time span of *Chlamydomonas* domestication (Pröschold, Harris, and Coleman 2005).

**Figure 6.**
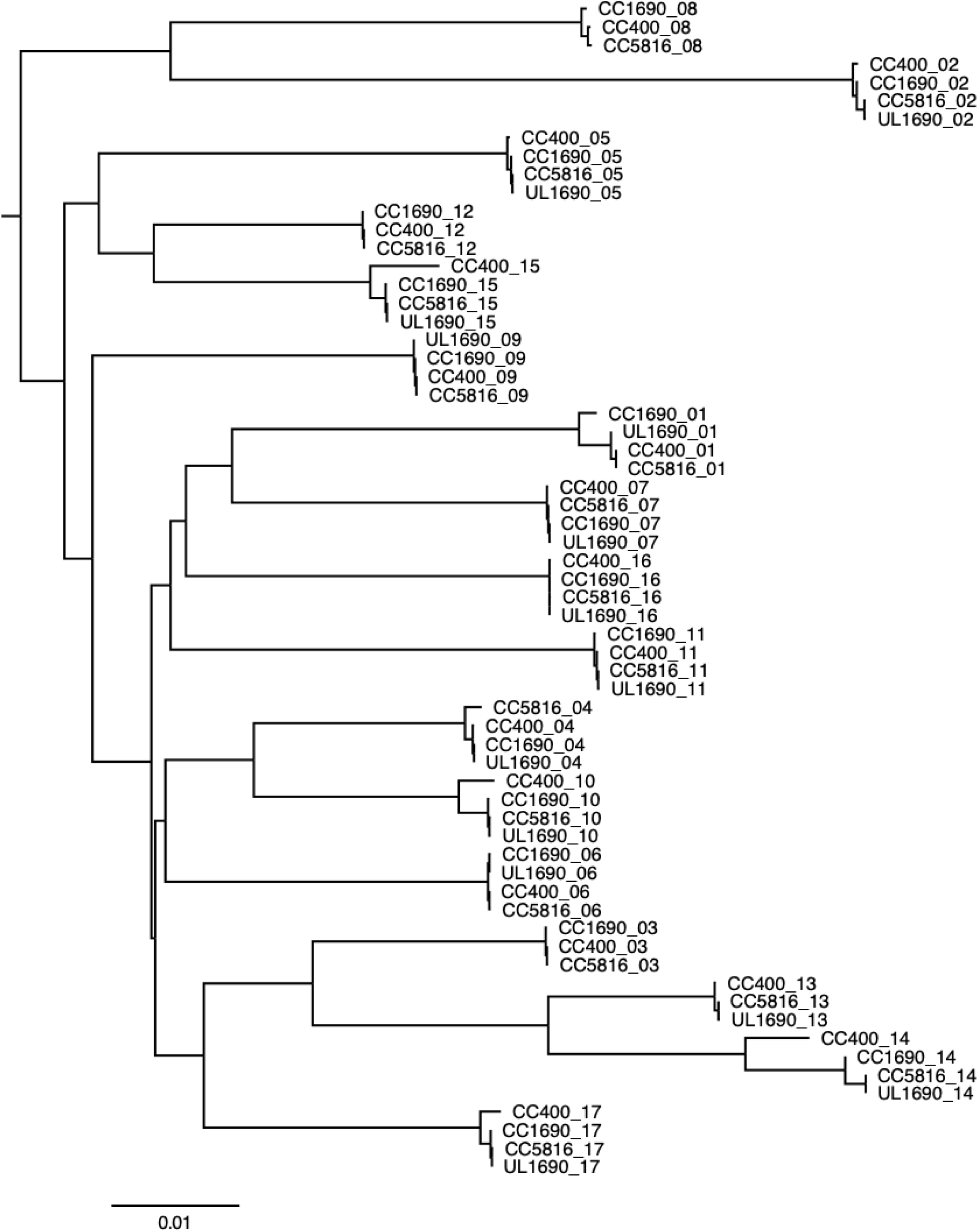
Alignment-free midpoint-rooted distance tree of all full-length centromere sequences from strains UL-1690, CC-1690, CC-400 and CC-5816 generated using Mash (Ondov et al. 2016) (see Materials and Methods for details). Labels show strain name followed by chromosome number. Relative distance scale is at the bottom.

## Discussion

CENH3 proteins and their association with centromeres have been studied in animals, fungi and land plants, but little work has been done on centromere proteins of green algae like Chlamydomonas (Maruyama et al. 2007) (Wang et al. 2024). Previous work defined centromeric regions in Chlamydomonas by genetic mapping (Kathir et al. 2003) and by the presence of ZeppL transposon repeats in assembled genomes (Craig et al. 2021, 2023). Here, we characterized two functionally overlapping CENH3 paralogs in *Chlamydomonas reinhardtii* and validated their expression and genomic localization at predicted centromere regions using CUT&Tag with custom CENH3 antibodies. We also introduce two newly assembled Chlamydomonas genomes and provide data suggesting that the transposition of ZeppL elements outside of the normal centromere domains can potentially result in neocentromere formation.

Most species encode only a single *CENH3/CENP-A* gene (Ishii et al. 2020; Elisafenko, Evtushenko, and Vershinin 2021) but in *Chlamydomonas reinhardtii* we found two. The two CENH3s exhibit significant divergence in their N-terminal regions (Figure 1a), suggesting that they did not arise from a very recent duplication event. Interestingly a search for *CENH3* orthologs in Volvocine algal relatives of *C. reinhardtii* identified only single genes even for its closest sequenced relative, *C. incerta* (Craig et al. 2021). Although this suggests a potentially recent duplication within the *C. reinhardtii* genome, the rapid evolution of CENH3 proteins makes this explanation difficult to distinguish from selective losses. However, the lack of *CENH3* duplicates in the nearest relatives of *C. reinhardtii* and most other species suggests that duplications of this gene are generally not long-lived.

In some species with two *CENH3* genes, one of the two paralogs appears to have a larger role than the other (Monen et al. 2005; Ishii et al. 2015, 2020). An extreme example is cowpea, where the localization patterns of the two different CENH3 paralogs generally overlap but only one of the two paralogs is essential for plant growth and reproduction (Ishii et al. 2020).

Unlike the case in cowpea, the two CENH3 paralogs in Chlamydomonas could both support apparently normal mitotic growth and meiotic segregation when the other paralog was absent making them seemingly redundant. However, we cannot rule out subtle phenotypes in our single knockout strains that would require more comprehensive analyses to detect.

One of the goals of this study was to prepare genome assemblies of sufficient quality to span the centromeres in two different *C. reinhardtii* strains, UL-1690 and CC-400. Most, but not all, the centromeres in these two strains were completely assembled. While we were carrying out our work, an entirely gapless assembly of strain CC-5816 was released (Payne et al. 2023). Each of these genomes are superior to the current assembly of the reference strain CC-4532 and include large repeat blocks absent from prior assemblies. Centromeres in all strains were marked by clusters of ZeppL elements that are 65-310 kb in length (Table S6). CENH3 CUT&Tag analysis confirmed that the functional centromeres lie within these repetitive regions (Figure 3). By measuring the distribution of uniquely mapped reads, we estimate that the functional domains are on the order of 50-105 kb (Table 1), considerably smaller than the estimated size of the centromeres in Arabidopsis, where the centromeric repeat arrays range from 10 to 22 Mb and the CENH3-enriched areas are roughly to ∼1-2 Mb (Wlodzimierz et al. 2023).

We observed evidence of active ZeppL transposition in laboratory strains since domestication (Figures S7-9, Data S2) as well as a very recent ZeppL transposition in strain UL-1690. The newly transposed ZeppL element on the end of chr2 is associated with what appears to be a new, secondary centromere (Figure 4b, Figure S6). Though we cannot assess CENH3 enrichment within the ZeppL element itself (due to the presence of three other ZeppL elements with nearly identical sequence (Data S2)), a total of ∼30 kb of unique sequence on either side of the ZeppL element is associated with CENH3 (Figure S6a), creating what appears to be a dicentric chromosome. While there are many known examples of newly formed centromeres (neocentromeres) (DeBose-Scarlett and Sullivan 2021; Comai, Maheshwari, and Marimuthu 2017), dicentric chromosomes should be unstable because the two centromeres will frequently migrate to opposite spindle poles and initiate cycles of breakage (McClintock 1941) that results in either the inactivation of one centromere, or the formation of two separate chromosomes (Stimpson, Matheny, and Sullivan 2012). The latter hypothesis is supported by a recent observation in maize, where activation of synthetic centromeres on maize chromosome 4 induced chromosome breakage, forming neo-chromosomes without disrupting gene expression, or normal plant development, and enabling stable inheritance (Zeng et al. 2025). Further work will be needed to determine if the neocentromere on chr2 is sufficiently large to independently interact with the spindle, or if chr2 is more unstable in UL-1690 than in other strains.

While it is theoretically possible that a neocentromere formed spontaneously on chromosome 2 after which ZeppL transposition occurred, the reverse seems more likely – that ZeppL transposed first and CENH3 was recruited over and around it. Most complex eukaryotes display a loose, epigenetic relationship between centromere sequence and CENH3 location (Comai, Maheshwari, and Marimuthu 2017). In contrast, ZeppL may have evolved sequence features that actively recruit CENH3. The best-known example of a genetic determination process affecting CENH3/CENP-A localization in multicellular eukaryotes comes from humans, where the major centromere repeat (the alpha satellite) contains a binding site for a protein known as CENP-B that facilitates CENP-A deposition (Otake et al. 2020). Similarly, in holocentric plants from the *Rhynchospora* genus, CENH3 is strictly correlated with the major centromere tandem repeat Tyba. Gain or loss of Tyba (over evolutionary time scales) results in a corresponding gain or loss of CENH3 binding (Hofstatter et al. 2022). Our data suggest that a single ∼8 kb ZeppL element may be sufficient to recruit CENH3. Due to the small size of its centromeres and a potentially simple route to new centromere formation driven by ZeppL transposon nucleation, Chlamydomonas may be an excellent model for engineering fully artificial plant chromosomes (Dawe 2024).

## Materials and Methods

### Chlamydomonas strains and growth conditions

Chlamydomonas strains CC-400 (cw15), CC-124, CC-125, CC-620, CC-621 were obtained from the Chlamydomonas Resource Center (https://www.chlamycollection.org/). UL-1690 is a mating type plus (*mt+*) strain, most likely derived from a cross between strains CC-1690 (21gr, *mt+*) and CC-1691 (6145c, *mt-*) that was mistakenly labeled as CC-1690 and has since been renamed UL-1690 (Umen Laboratory 1690). All strains were maintained on Tris-acetate-phosphate (TAP) + 1.5% BD Difco agar (Thermo Fisher, Cat# DF0812-07-1) plates. For asynchronous growth, cells were cultured at 25°C in TAP liquid (https://www.chlamycollection.org/methods/media-recipes/tap-and-tris-minimal/) with illumination from red (625 nm, 100 µE) and blue (465 nm, 100 µE) LEDs and aeration from bubbling with air. For synchronous growth, cells were cultured at 25°C in Sueoka’s high salt medium (HSM) (Sueoka 1960) under a 12hr dark: 12hr light diurnal cycle with illumination from red (625 nm, 150 µE) and blue (465 nm, 150 µE) LEDs and aeration from bubbling with 1% CO2. Gamete generation, mating, and zygote germination were performed following standard protocols (Harris 1989). Segregation analysis was done with random meiotic progeny sub-cloned after bulk germination.

### Antibody generation

Rabbit polyclonal antibodies were prepared, and affinity purified by Pacific Immunology, Ramona, CA. The peptides were designed to uniquely identify CENH3 proteins but not canonical histone H3. To generate antibodies unique to CrCENH3.1, we used a peptide matching amino acids 15-39 (KAQAEAATPTKSKRPSGAAATPTRG-Cys), to generate antibodies unique to CrCENH3.2 we used a peptide matching amino acids 15-40 (EAAAQRSTRAGEAVGSPARGGTPARS-Cys) and to generate antibodies that would identify both CrCENH3.1 and CrCENH3.2 we used a peptide matching amino acids 8-16 (PARPGRKA-Cys, though note that the lysine at position 15 is only found on CrCENH3.1).

### Nuclear protein enrichment and extraction

The nuclear protein enrichment protocol was adapted and simplified from (Rommelfanger et al. 2021). Chlamydomonas cultures were grown synchronously (described above) and maintained at a cell concentration between 5×10^5^ cells/mL to 1×10^6^ cells/mL. 5×10^8^ daughter cells (light phase ZT=1) per cell culture were harvested by centrifugation at 3,000 g for 5 min with 0.005% Tween20. Pellets were flash frozen in liquid nitrogen, then thawed on ice in 1mL nuclear isolation buffer (NIB, 25mM HEPES pH7.0, 20mM KCl, 20mM MgCl2, 0.6M sucrose, 10% glycerol, 1mM PMSF, 5mM DTT, 10mM sodium butyrate, 1X Roche cOmplete Protease Inhibitor Cocktail, Roche 04693116001) + 5% Triton X-100. The thawed pellets were flash frozen in liquid nitrogen again, then subject to tissue grinding twice (frequency 30 Hz, duration 90 seconds, baskets chilled with liquid nitrogen) with three 3mm glass beads per sample in 1.5mL Eppendorf tubes using a Qiagen TissueLyzer II. The macerated mixture was then thawed and pelleted by centrifugation at 1,000 g for 60 min at 4°C. Pellets were washed twice using NIB and recollected by centrifugation 1,000 g for 30 min at 4°C. Nuclear-enriched pellets were resuspended in lysis buffer (1xPBS pH 7.4, 1x Roche protease Inhibitor, 500 mM PMSF) to a final concentration equivalent of ∼ 5×10^9^ cells/mL. The suspensions were solubilized using a Covaris ultrasonicator (peak power 150, duty factor 150, cycle 200, treatment 120 sec, 4°C) to generate protein lysates. Protein lysates were mixed 5:1 with 6X SDS protein loading buffer and boiled for 10 min, then used either directly in immunoblotting or saved for later uses at -20°C.

### Immunoblotting

Protein lysates in 1X loading buffer were further cleared by centrifugation at 12,000 g for 10 min. Supernatants were loaded onto 15% tricine sodium dodecyl sulfate-polyacrylamide (SDS-PAGE) gels for separation, then wet-transferred to polyvinylidene fluoride membranes for 1hr at 50V (Blancher and Jones 2001). Membranes were blocked in PBS containing 5% nonfat dry milk, incubated with primary antibody (anti-CENH3.1, 1:1,000; anti-CENH3.2, 1:500; anti-CENH3-conserved, 1:500) added to PBS containing 3% non-fat dry milk overnight at 4°C. Membranes were then washed in PBS containing 0.1% Tween20 for 3 × 10 min. Secondary antibodies coupled to horseradish peroxidase (goat anti-rabbit 1:10,000, Thermo Scientific) were added to PBS containing 3% non-fat dry milk. Membranes were incubated with a secondary antibody for 1 hr at room temperature, then washed in PBS containing 0.1% Tween20 for 3 × 10 min. Chemiluminescent detection was done using a Bio-Rad Chemidoc XRS+ Imaging System.

### Immunofluorescence microscopy

Wild type UL-1690 cells (∼10^7^ cells) were grown asynchronously in TAP (described above) and maintained at a cell concentration between 5×10^5^ cells/mL to 1×10^6^ cells/mL. 5×10^8^ cells per cell culture were harvested by centrifugation at 3,000 g for 5 min with 0.005% Tween20 and resuspended in 240uL fixation buffer (2% paraformaldehyde in PBSP (1x PBS pH7.4, 1 mM DTT, 1x Roche protease inhibitor)) and left for 30 min on ice. Fixed cells were extracted in cold methanol 3 x 10 min at -20°C and rehydrated in 1mL PBSP for 30 min on ice. Fixed cells were blocked for 30 min in 1mL blocking solution I (5% BSA and 1% cold water fish gelatin in PBSP) and 30 min in 1mL blocking solution II (10% goat serum, 90% blocking solution I) at room temperature. Cells were then incubated overnight with one of three primary antibodies (anti-CENH3.1, 1:200; anti-CENH3.2, 1:50; anti-CENH3-conserved, 1:50) in 20% blocking solution I total volume 50uL at 4°C, then washed 3 x 10 min in 1% blocking solution I at room temperature. Cells were incubated with 1:1,000 Alexa Fluor 488 conjugated goat anti-mouse IgG (Thermo Fisher) in 20% blocking solution I total volume 100uL for 1 hr at 4°C and then incubated with 4’,6-diamidino-2-phenylindole, dihydrochloride (DAPI) at a final concentration of 5ug/mL at room temperature for 5 min. Cells were washed in 1 x PBS for 3 x 10 min, then mounted for microscopy in 9:1 Mowiol: 0.1% 1, 4-phenylenediamine (PPD). Cells were imaged with a Zeiss Elyra7 Super-Resolution microscope using a 63X oil lens with 405nm (50mW, 4.5%), 488nm (500mW, 4.0%), and 561nm (500mW, 4.0%) diode laser lines, with filter settings 561 TV1:BP 570-620 + LP 655, 488 TV2: BP 420-480 + BP 495-550, and 405 TV2: BP 420-480 +BP 495-550. 3D LEAP MODE was used to capture volumes of ∼ 3 µm thickness in ∼ 100nm increments. The image stacks were further processed with Zeiss Zen Black image analysis software using the dual iterative SIM (SIM²) algorithm (Löschberger et al. 2021), 3D leap processing mode (default parameters, sharpness 3.0) for all channels, and exported as a single maximum projection images.

### Chlamydomonas transformation

Chlamydomonas cultures were grown to 10^6^ cells/mL were grown under the asynchronous condition (described above) and harvested by centrifugation at 3,000g for 5 min with 0.005% Tween20. Cell pellets were resuspended and washed in TAP + 40 mM sucrose to a final concentration of 2×10^8^ cells/mL. 125 µL cell suspension (∼ 2.5×10^7^ cells) was mixed with ∼ 500 ng plasmid and transferred into a 2-mm cuvette. Electroporation was performed using a NepaGene Electroporator with poring pulse (250 V initial, 2 pulses, 8 ms each, 40% decay, 50 ms interval) and transfer pulse (20 V initial, 5 pulses with alternating polarity, 50 ms each, 40% decay, 50 ms interval). Cells were then transferred into 5 mL TAP+40 mM sucrose and recovered under dim light at room temperature overnight before plating on antibiotic selection plates as specified below.

### CRISPR-Cas9 mediated mutagenesis of *CENH3.1* and *CENH3.2*

gRNAs were designed using the CRISPR RGEN tool (http://www.rgenome.net/cas-designer/). The chosen gRNA target sites are listed in Table S1. *In vitro* transcription of sgRNA was performed following (Yu et al. 2017). 5 μg (∼30 pmol) Cas9 Nuclease (Alt-R™ S.p. Cas9 Nuclease V3, IDT Cat# 1081059) was incubated with 5,000ng (∼120 pmol) purified sgRNA in 3uL IDT RNA duplex buffer at 37°C for 30 minutes to assemble RNP complex for one transformation.

CRISPR-Cas9 targeted insertional mutagenesis was performed following (Greiner et al. 2017; Picariello et al. 2020) with minor modifications: Cultures of strain UL-1690 were grown to 2×10^6^ cells/mL asynchronously in TAP (described above) and harvested by centrifugation at 2,000 g for 5 mins. Pellets were resuspended in 6mL autolysin (Findinier 2023) and incubated at 33°C for 30 mins, then heat shocked at 40 °C for another 30 mins. After heat shock, cells were collected by centrifugation at 2,000 g for 2 mins, then resuspended and washed in TAP + 40mM sucrose to a final cell concentration 2×10^8^ cells/mL. 125 µL cell suspension (∼2.5×10^7^ cells) were mixed with RNP complex and 500 ng Chlamydomonas paromomycin marker (linear PCR product using M13 primer sets from pKS-aphVIII-lox (Sizova, Fuhrmann, and Hegemann 2001; Heitzer and Zschoernig 2007)). The RNP and donor DNA were delivered into the cells by electroporation as described above.

Individual transformants were selected on TAP + 1.5% BD agar with 20 μg/mL paromomycin on a light shelf (50uE white light) at room temperature for 6 days. Transformants were then picked and re-grown individually in 96 well plates with 200uL TAP medium for 3 days. Crude DNA preparation for genotyping followed (Zamora, Feldman, and Marshall 2004).

Transformants were screened by genotyping for insertion of the paromomycin marker at the target site. Genotyping oligos and amplification conditions are listed in Table S2. Genotyping PCR reactions were performed in a 10 μL volume containing 200 μM of each dNTP, 0.4 μM of each primer, 3% DMSO, 1 μL of homemade recombinant Taq DNA polymerase, and 1 μL of crude DNA preparation. Homemade Taq buffer final concentrations: 10mM Tris-HCl pH8.4, 50mM KCl, 1.5mM MgCl2, 0.08% NP40, and 0.4ug/uL BSA. Loss of CENH3 expression was further confirmed by immunoblotting (see above).

### Transgenic rescue of *cenh3* mutants

A ∼4.7kb fragment containing the full-length genomic region of *CENH3.1* and its upstream region (∼1.8kb upstream of the transcription start site) was amplified with 2x KOD Hot Start Master Mix (Sigma-Aldrich Cat# 71842) from genomic UL-1690 DNA using primers CENH3.1 comp F1/ CENH3.1 comp R1 (Table S2). Vector backbone was amplified with 2x KOD Hot Start Master Mix from pHsp70A/RbcS2-cgLuc (Heitzer and Zschoernig 2007) using primers CENH3.1 IO OH R1/ CENH3.1 IO OH F1 (Table S2). The amplified vector fragment was ligated with the amplified *CENH3.1* genomic region using a Takara In-Fusion kit (In-Fusion Snap Assembly Master Mix, Cat # 638947) to generate *CENH3.1* rescue construct pCENH3.1.

A ∼4.6kb fragment containing the full-length genomic region of *CENH3.2* with ∼1.5kb upstream of 5’UTR was amplified with 2x KOD Hot Start Master Mix (Sigma-Aldrich Cat# 71842) from genomic CC-1690 DNA using primers CENH3.2 comp F2/ CENH3.2 comp R2 (Table S2). Backbone vector pHsp70A/RbcS2-cgLuc (Heitzer and Zschoernig 2007) was amplified with 2x KOD Hot Start Master Mix using primers CENH3.2g IO OH R2/ CENH3.2g IO OH F2 (Table S2). The amplified vector fragment was ligated to the genomic fragment of *CENH3.2* using a Takara In-Fusion kit to generate *CENH3.2* rescue construct pCENH3.2 (constructs maps in Figure S2e).

Either pCENH3.1 or pCENH3.2 (500ng/transformation) were co-transformed with pKS-aphVII-lox (50ng/transformation) (Heitzer and Zschoernig 2007; Berthold, Schmitt, and Mages 2002) into Chlamydomonas *cenh3.1* or *cenh3.2* mutants (Methods see above) by electroporation. Transformants were selected on TAP + 1.5% BD agar with 30 μg/mL hygromycin. Individual transformants were picked into 96 well plates and screened for the presence of *CENH3* transgenes by PCR genotyping (Table S2). Crude DNA preparation for genotyping followed (Zamora, Feldman, and Marshall 2004). All genotyping oligos and amplification conditions are listed in Table S2. Genotyping PCR reactions were performed in a 10 μL volume containing 200 μM of each dNTP, 0.4 μM of each primer, 3% DMSO, 1 μL of homemade recombinant Taq DNA polymerase, and 1 μL of crude DNA preparation. Homemade Taq buffer final concentrations: 10mM Tris-HCl pH8.4, 50mM KCl, 1.5mM MgCl2, 0.08% NP40, and 0.4ug/uL BSA.

### PacBio HiFi sequencing and Genome assemblies

Chlamydomonas cultures were grown under asynchronously in TAP (described above) and harvested by centrifugation at 3,000g for 5 min with 0.005% Tween20. 4g of wet weight pellets were washed twice in TAP, flash frozen in liquid nitrogen, and sent to Arizona Genomics Institute (AGI) for high molecular weight DNA preparation and HiFi PacBio sequencing. High molecular weight DNA was extracted from cell pellets using the protocol described in (Gontero et al. 2017). DNA was sheared to 10-30 kb using Megaruptor 3 (Diagenode) and purified with PB Beads (Pacbio, Menlo Park, CA). Sequencing libraries were constructed using SMRTbell Express Template Prep kit 3.0 (Pacbio, Menlo Park, CA) and size selected to 10-25 kb on a Blue Pippin (Sage Science). The libraries were prepared for sequencing with PacBio Sequel II Sequencing kit 2.0 for HiFi library, loaded to 8M SMRT cells, and sequenced in CCS mode in the Sequel II instrument for 30 hours. Approximately 82x (UL-1690) and 89x (CC-400) HiFi data were assembled to primary contigs using Hifiasm (version 0.19.6) (Cheng et al. 2021) (parameter: -l 0). The N50 values after assembly were 15818 Kb for UL-1690 and 14254 Kb for CC-400. To generate chromosome-level sequence scaffolds, RagTag (version 2.1.0) (Alonge et al. 2022) “correct” and “scaffold” with default parameters were used to correct potential misassembled contigs and orient contigs relative to the published CC-5816 genome (Payne et al. 2023).

Centromere variations observed between the sequenced strains and the reference strain are genuine and not due to assembly errors. First, assembly errors often occur near gap regions, particularly in areas with limited long-read coverage, repetitive sequences, or structural variations. These errors can be more pronounced at contig edges before pseudochromosome construction due to incomplete sequence information and misassembles. However, in our sequenced strains, centromere regions contain fewer gaps—three gaps in UL-1690 (chromosomes 3, 8, and 12) and no gaps in CC-400—suggesting that gaps have minimal impact on centromere contiguity. Second, long reads spanning the edges of centromere regions (ZeppL enriched sequences of UL-1690, CC-1690, and CC-400) and the reference assembly (CC-5816) were identified in our genome assemblies, further supporting the accuracy of the genome assemblies among these centromere variations.

### Gene annotation

UL-1690 gene models were annotated based on the gff3 file of the reference strain CC-4532 (Goodstein et al. 2012). Liftoff (version 1.6.3) (Shumate and Salzberg 2021) was used to map annotations from CC-4532 to the UL-1690 assembly (parameter: -exclude_partial - polish). The longest isoforms for each gene were kept using AGAT for gene model visualization in IGV and gene density calculation (Dainat, 2024). Gene density was calculated by counting the gene number per 5 kb window size.

### CENH3 CUT&Tag

Active Motif CUT&Tag-IT Assay Kit (Rabbit antibody specific, Cat# 53160) was used according to the manufacturer’s instructions with specific adaptations we developed for Chlamydomonas cells. UL-1690 was synchronized as described above and ∼ 5×10^7^ daughter cells (light phase ZT=1) per reaction were harvested by centrifugation at 3,000g for 5 min with 0.005% Tween20. Harvested cells were subject to autolysin treatment (described above) to remove their walls prior to incubation with concanavalin-A coated magnetic beads supplied in the kit. The remaining steps followed the manufacturer’s instructions but using 1:70 5% digitonin instead of the suggested 1:50 5% digitonin. 1 ug primary antibody (anti-CENH3.1, anti-CENH3.2, anti-CENH3-conserved), or rabbit IgG Control (Invitrogen, Rabbit IgG Isotype, Cat# 10500C) in a 50uL reaction were used for each procedure.

All primary libraries were amplified with 14 cycles of PCR. After combining libraries together, they were purified using a double-sided size selection and bead clean up with Mag-Bind TotalPure NGS (Omega Bio-Tek, M1378-00). In the first size selection, large DNA molecules were removed using a 0.5:1 volume ratio of beads to DNA, then in the second size selection, small DNA molecules were removed using a 1:1 volume ratio of beads to DNA. Illumina sequencing was carried out with paired-end, 151-nt read lengths.

### CUT&Tag data analysis

Centromere regions were determined by identifying CENH3 CUT&Tag enriched regions. The adapter sequences of all Illumina reads were trimmed by TrimGlore (version 0.6.7) (https://github.com/FelixKrueger/TrimGalore/releases) (parameter: --fastqc –paired --gzip). All trimmed reads were aligned to the genome sequence using Bowtie2 (version 2.4.5) (Langmead and Salzberg 2012) (parameter: --end-to-end --very-sensitive --no-mixed --no-discordant -- phred33 -I 10 -X 700). Two versions of the alignment BAM file (multiple plus unique mapping and unique mapping only) were obtained for each of the four datasets from four antibodies using Samtools (version 1.16.1) (Danecek et al. 2021). Multiple plus unique mapping was used to estimate the maximum CENH3 CUT&Tag enriched regions. Unique mapping, which was the output after removing all reads with MAPQ scores lower than 20, was used to estimate the minimum CENH3 CUT&Tag enriched regions. All alignment BAM files, including IgG controls, were converted to bedgraph files using Bedtools “genomecov” function (version 2.30.0) (Quinlan and Hall 2010), and then uploaded to an Integrated Genome Viewer browser for data visualization (Robinson et al. 2011). All BAM files were converted to bed files using Bedtools “bamtobed” function. The read depth for each 5 kb bin of the UL-1690 HiFi genome was calculated with Bedtools “coverage” function (version 2.30.0) (Quinlan and Hall 2010). To eliminate noisy or false CENH3 read enrichment peaks, genomic loci with the counts of control IgG reads lower than 10 were corrected to 20 manually. The normalized CENH3 and IgG CUT&Tag read depth was calculated by dividing CUT&Tag read counts by total read counts in the same 5kb bin. The fold enrichment of CENH3 reads was calculated by dividing the normalized CENH3 CUT&Tag read depth by the normalized IgG CUT&Tag read depth. CENH3 CUT&Tag enriched regions were plotted using the karyoploteR package (Gel and Serra 2017) in R.

### Centromere size estimation

Multiple plus unique mapping CENH3.1 CUT&Tag reads were used to estimate the upper centromere sizes, and unique mapping reads alone were used to estimate lower centromere sizes for each chromosome in the UL-1690 genome. For the upper centromere size estimation, genomic loci with fold enrichment > 5.5 were regarded as stringent CENH3 peaks and then were merged (intervals between adjacent bins were < 30 kb) using Bedtools (version 2.30.0) “merge” function (Quinlan and Hall 2010). The same method was used to calculate the lower centromere sizes, except that genomic positions with fold enrichment < 10 were considered as stringent CENH3 peaks. 18 merged CENH3 CUT&Tag enriched regions larger than 10 kb that overlap with ZeppL-enriched regions were identified as centromere positions (17 genetic centromeres + 1 potential neo-centromere on chr2).

### Tandem repeats and ZeppL enriched regions identification

The CC-5816 and CC-1690 *Chlamydomonas reinhardtii* genome assemblies were downloaded from GenBank accession number GCA_026108075.1 (Payne et al. 2023) and JABWPN000000000 (O’Donnell et al., 2020). The CC-4532 PacBio assembly (Craig et al. 2023) was downloaded from Phytozome. The coordinates of ZeppL were identified by BLAST (Altschul et al. 1990) search using the published ZeppL sequence (Craig et al. 2023) in UL-1690, CC-400, CC-4532, CC-1690 and CC-5816 genomes. For tandem repeats regions, genomes were annotated using Tandem repeats finder with parameter: 2 7 7 80 10 50 2000 -d -h (Benson 1999). The unique tandem repeats with copies lower than 400 and the period size lower than 125 were filtered. For ZeppL enriched regions, Bedtools “merge” was used to merge nearby ZeppL sequences if the intervals of their closest coordinates were < 40 kb. ZeppL enriched regions overlapping with CENH3 peaks are shown in Tables S5 and S6.

### Dot plot comparisons of centromere regions

Centromere regions defined based on ZeppL enrichment (see above) were extracted from UL-1690, CC-400, CC-1690 and CC-5816 genomes individually. CC-4532 centromere regions were excluded due to too many assembly gaps in its centromere regions. All centromere regions from UL-1690, CC-400, CC-1690 and CC-5816 (UL-1690 contains 18 regions = 17 centromeres + 1 neocentromere) were divided into bin windows of 500bp and compared to the complete centromere regions from CC-5816 using BLAST (Altschul et al. 1990). The output of the BLAST search was visualized using dot plots (Mohanty, Sahoo, and Mishra 2022) of the bit score from BLAST.

### Centromere comparisons between chromosomes and strains

From the 68 genetic centromere regions described above, 64 were gapless and used for alignment free comparison using Mash software (Ondov et al. 2016). A k-mer size of 9 was chosen empirically after sampling k=7 to k=15 and finding very similar results for k=8 and above. A sketch size of 10^6 was used for k-mer sampling but smaller sketch sizes down to 10^3 yielded similar results. The output distance matrix was converted to a distance tree using the T-REX web server (Boc, Diallo, and Makarenkov 2012) and the tree was visualized and formatted using FigTree v1.4.4 (https://github.com/rambaut/figtree/releases/tag/v1.4.4).

### Phylogeny of full length or near full length ZeppL elements

All ZeppL elements larger than 8kb were extracted from the UL-1690, CC-400, CC-1690 and CC-5816 genomes producing a set of 49 full length or near full length ZeppL elements among these strains (Data S2). The sequences were aligned with MAFFT and used for construction of a neighbor joining tree in MEGA11 (Tamura, Stecher, and Kumar 2021) set with the Tamura 3-parameter model + gamma (0.05) and 500 bootstrap replicates.

## Data availability

All sequencing data generated in this study have been deposited in the NCBI BioProject database under accession number PRJNA1060758, including PacBio raw reads and CUT&Tag Illumina reads. The genome assemblies for strains UL-1690 (GCA_047496495.1) and CC-400 (GCA_047496505.1) are also available within the same BioProject. All scripts used for data analysis and visualization are publicly accessible on GitHub at https://github.com/dawelab/Chlamydomonas_centromeres.

## Acknowledgements

We thank Dong won Kim, Brooke Harris, Zach Jaudes, Ashley Cloud, Kayleigh Kreher, and Chris Reynolds for laboratory support. We thank Dr. Rebecca Bart, Ke Ke, and Marisa Yoder for the training on Bio-Rad Image-Lab software and Chemi Doc XRS+ Imaging System. We thank Dr. Rebecca Bart and Ke Ke for the training on a Qiagen TissueLyzer. We thank Dr. Kirk Czymmek and Dr. Anastasiya Klebanovych for the guidance and assistance on microscopy. This work was funded by NSF PGRP IOS 2151105 to Dr. Kelly Dawe and Dr. James Umen and supported through NSF award MRI 2018962 to the Donald Danforth Plant Science Center for acquisition of a super resolution microscope.

## Supporting information

### Supporting figures

**Figure S1.**
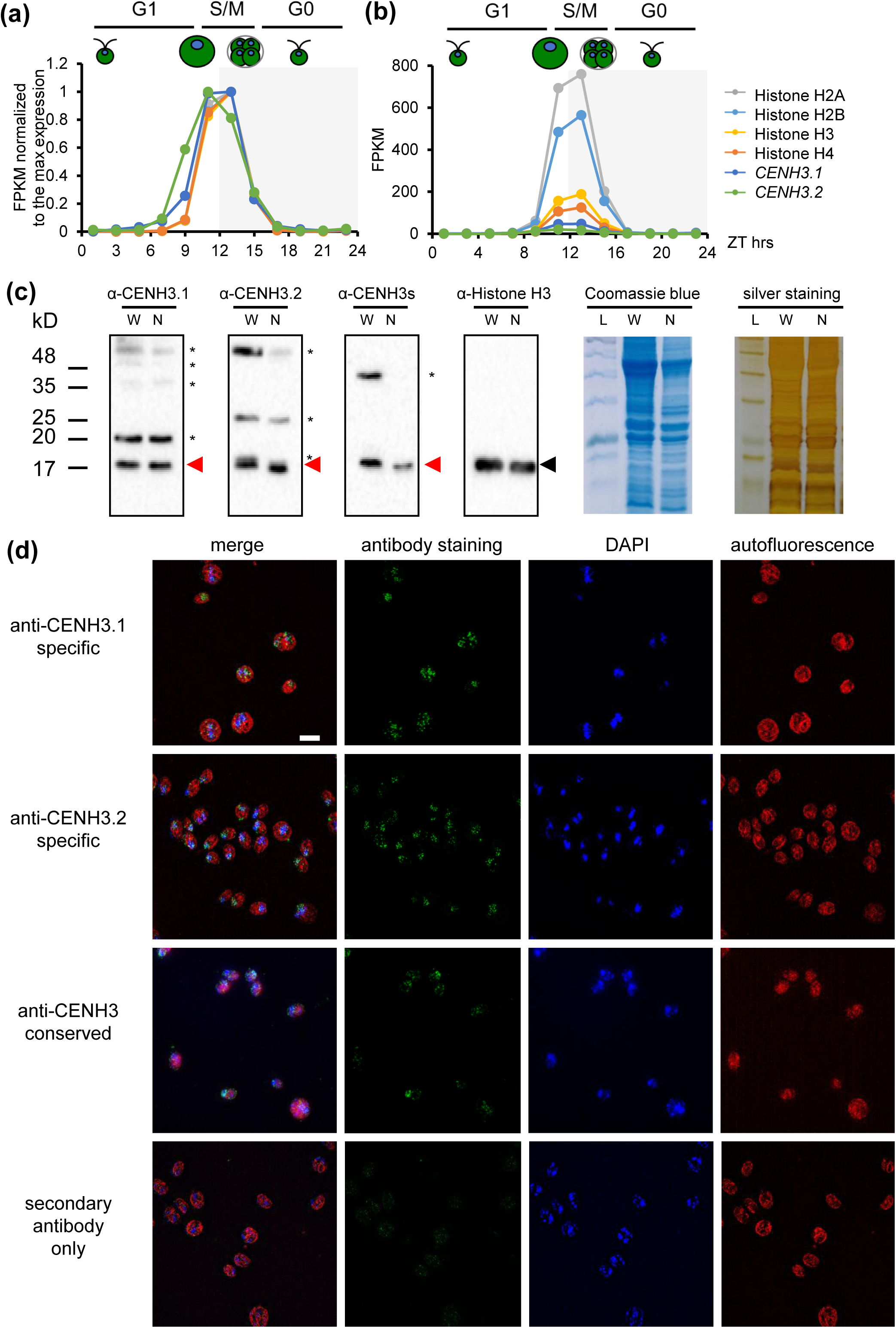
(a-b) Expression patterns of *CENH3.1* (Cre16.g661450) and *CENH3.2* (Cre02.g104800) with representative replication-dependent histones H3 (Cre17.g713950), H4 (Cre17.g714000), H2A (Cre17.g714100), and H2B (Cre17.g714050) for comparison. Data from previously published expression estimates of synchronous diurnal wild-type cultures (Strenkert et al., 2022) were used to generate the figures. Light phase and dark phase are indicated by white and gray backgrounds, respectively. DNA replication and cell division (S/M phase) occur at the beginning of the dark period concurrent with expression of replication dependent histones. **(a)** The maximum expression for each gene was normalized to 1. **(b)** FPKM (Fragments Per Kilobase of transcript per Million mapped reads) plots without internal normalization show relative expression magnitudes of each gene. **(c)** Full immunoblots of SDS-PAGE separated whole cell protein lysates (W) and nuclear protein enriched protein lysates (N) of wild type using α-CENH3.1-specific, α-CENH3.2-specific, α-CENH3-conserved, or α-histone H3 (internal loading control). The black arrowhead shows the position of histone H3. The red arrowheads show the position of CENH3 proteins. Asterisks mark cross-reacting non-specific antigens. Coomassie Blue and Silver staining of parallel gels serve as loading controls for total input and protein profiles. L denotes the ladder. **(d)** Immunofluorescence microscopy images of strain UL-1690. Cells were immunostained with CENH3.1-specific, CENH3.2-specific, CENH3-conserved, or IgG (negative control) antibodies and imaged as in Figure 1c. Scale bar = 20 μm.

**Figure S2.**
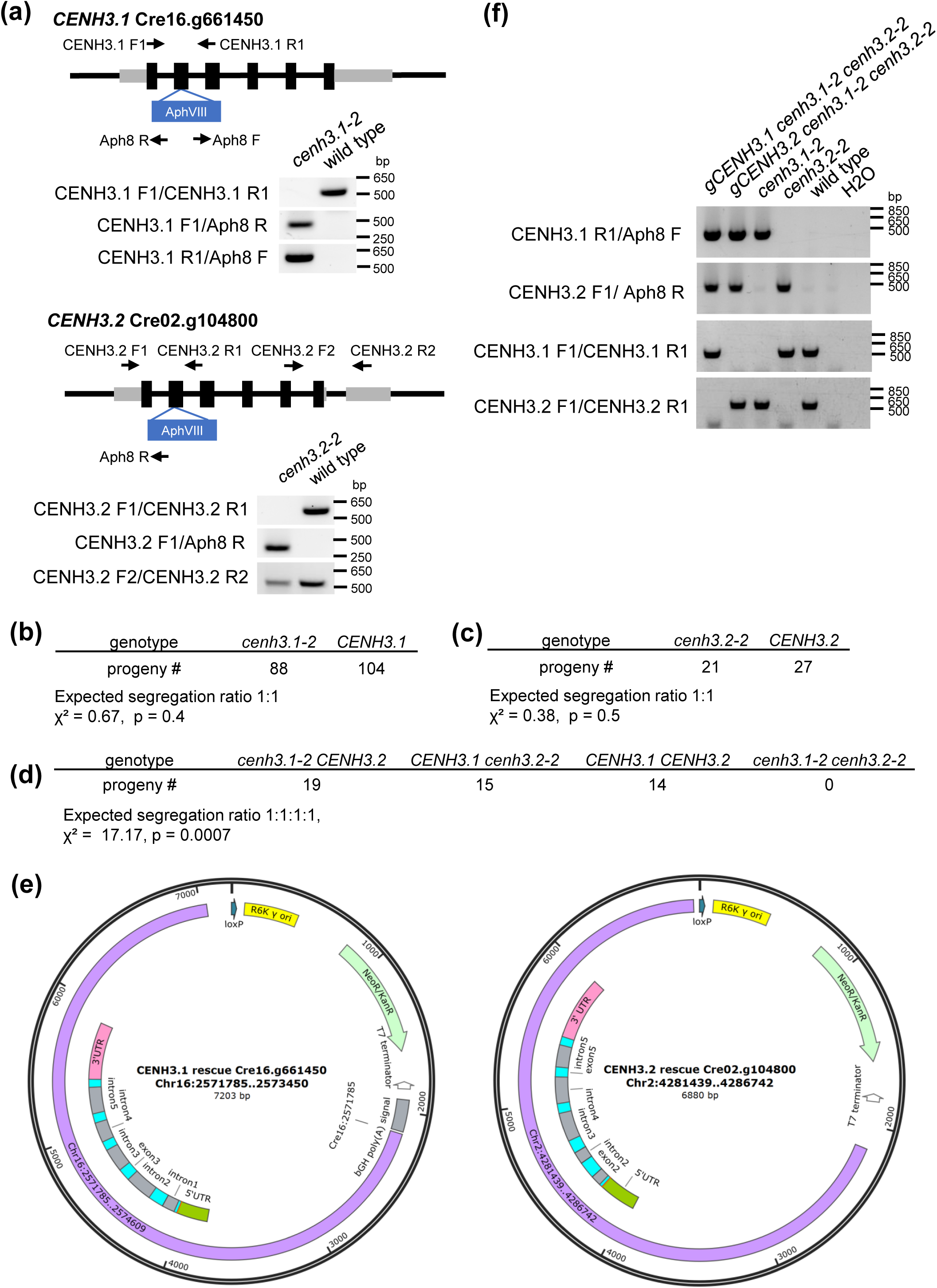
(a) Genotyping *AphVIII* insertions in *cenh3.1 and cenh3.2* mutants. Schematic of *CENH3.1* and *CENH3.2* loci with location of inserted paromomycin resistance markers (*AphVIII* in blue) and locations of genotyping PCR primer binding sites indicated by black arrows. *cenh3.1-2* was created by insertion of AphVIII in the 2^nd^ exon of *CENH3.1* and both borders could be amplified (primer sets CENH3.1 F1/Aph8 R and CENH3.1 R1/Aph8 F). *cenh3.2-2* was created by an insertion of *AphVIII* in the 2^nd^ exon. The left junction of *cenh3.2-2* was successfully amplified (primer set CENH3.2 F1/Aph8 R), but not the right junction. The 5^th^ exon and 3’UTR region of *cenh3.2-2* could be amplified indicating that there is not a deletion extending outside of the locus (primer set CENH3.2 F2/CENH3.2 R2). **(b-d)** Meiotic segregation ratios of progeny from **(b)** *cenh3.1-2* crossed to wild-type strain (CC-1691/6145c), **(c)** *cenh3.2-2* crossed to wild-type strain (CC-1691/6145c), and **(d)** *cenh3.1-2* outcrossed with *cenh3.2-2* mutant. Chi squared tests of each cross are shown below the segregation data. **(e)** Maps of constructs used to rescue *cenh3.1-2* and *cenh3.2-2* mutants with native genomic *CENH3.1* or *CENH3.2* genes respectively. **(f)** Genotyping for indicated strains to detect presence/absence of mutant and wild type *CENH3.1* and *CENH3.2* alleles in rescued *cenh3.1-2 cenh3.2-2* double mutant strains (the first two lanes).

**Figure S3.**
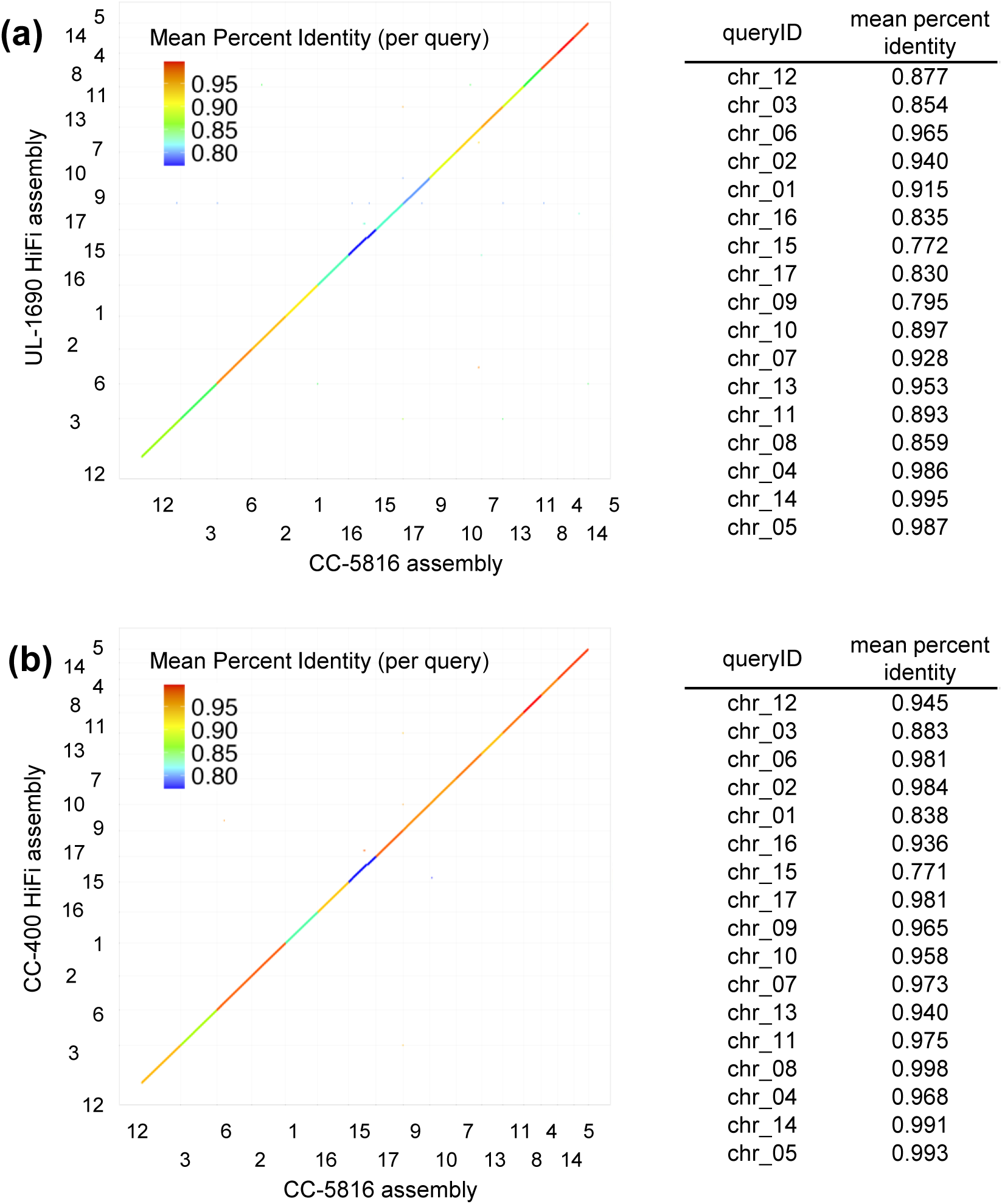
Comparison of new Chlamydomonas genome assemblies for UL-1690 and CC-400 against gap-free reference assembly CC-5816. Plot of the 17 largest scaffolds (corresponding to 17 chromosomes) from **(a)** UL-1690 HiFi, and **(b)** CC-400 HiFi aligned against the CC-5816 assembly using dotPlotly (https://github.com/tpoorten/dotPlotly/blob/master/pafCoordsDotPlotly.R). Inset table shows the mean percent identity score for each chromosome.

**Figure S4.**
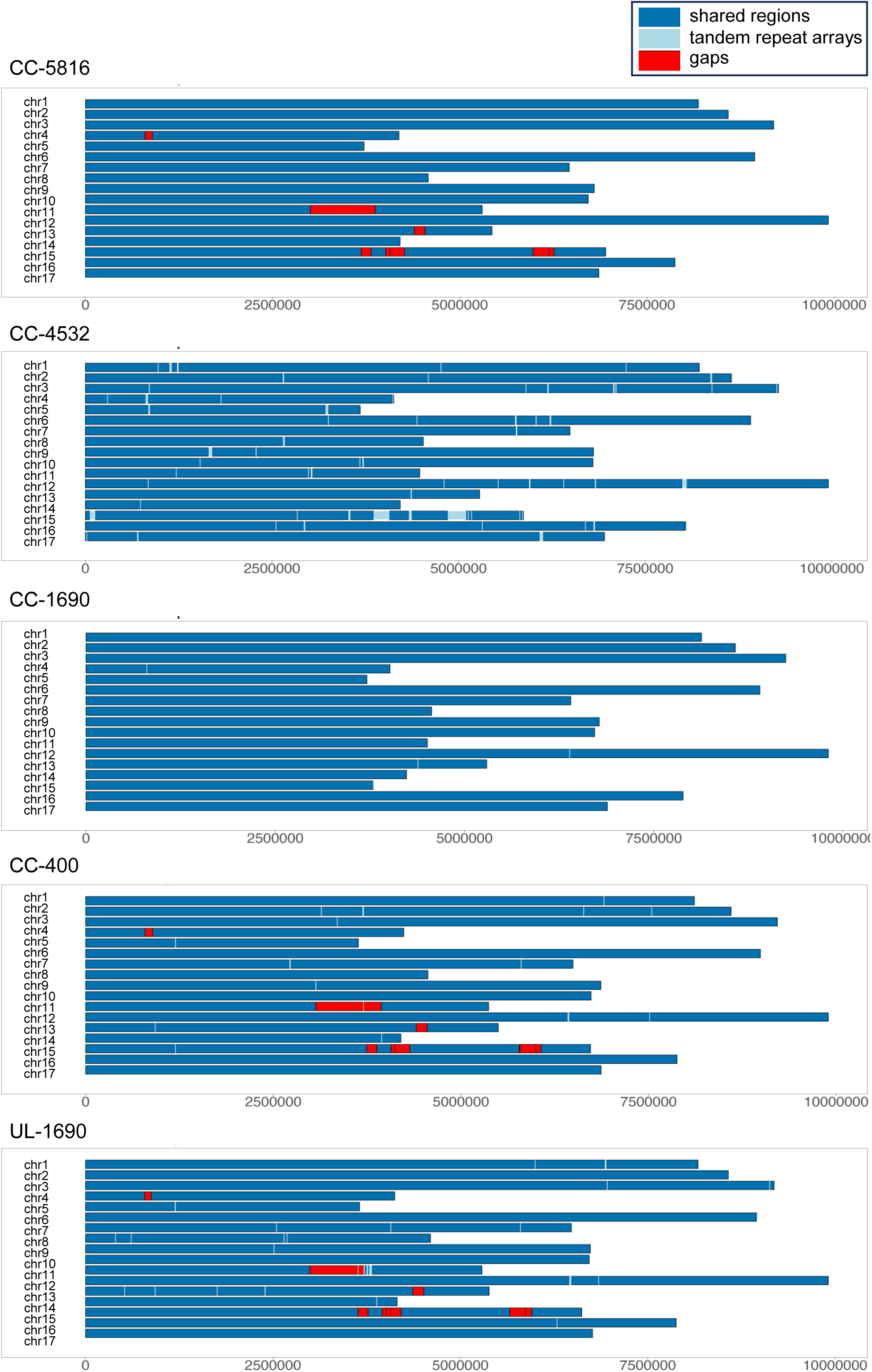
Comparison of five Chlamydomonas genome assemblies with regions common to all assemblies shown in dark blue and locations of long tandem repeat arrays and gaps shown in light blue and red, respectively. The x axis indicates chromosome lengths in bp.

**Figure S5.**
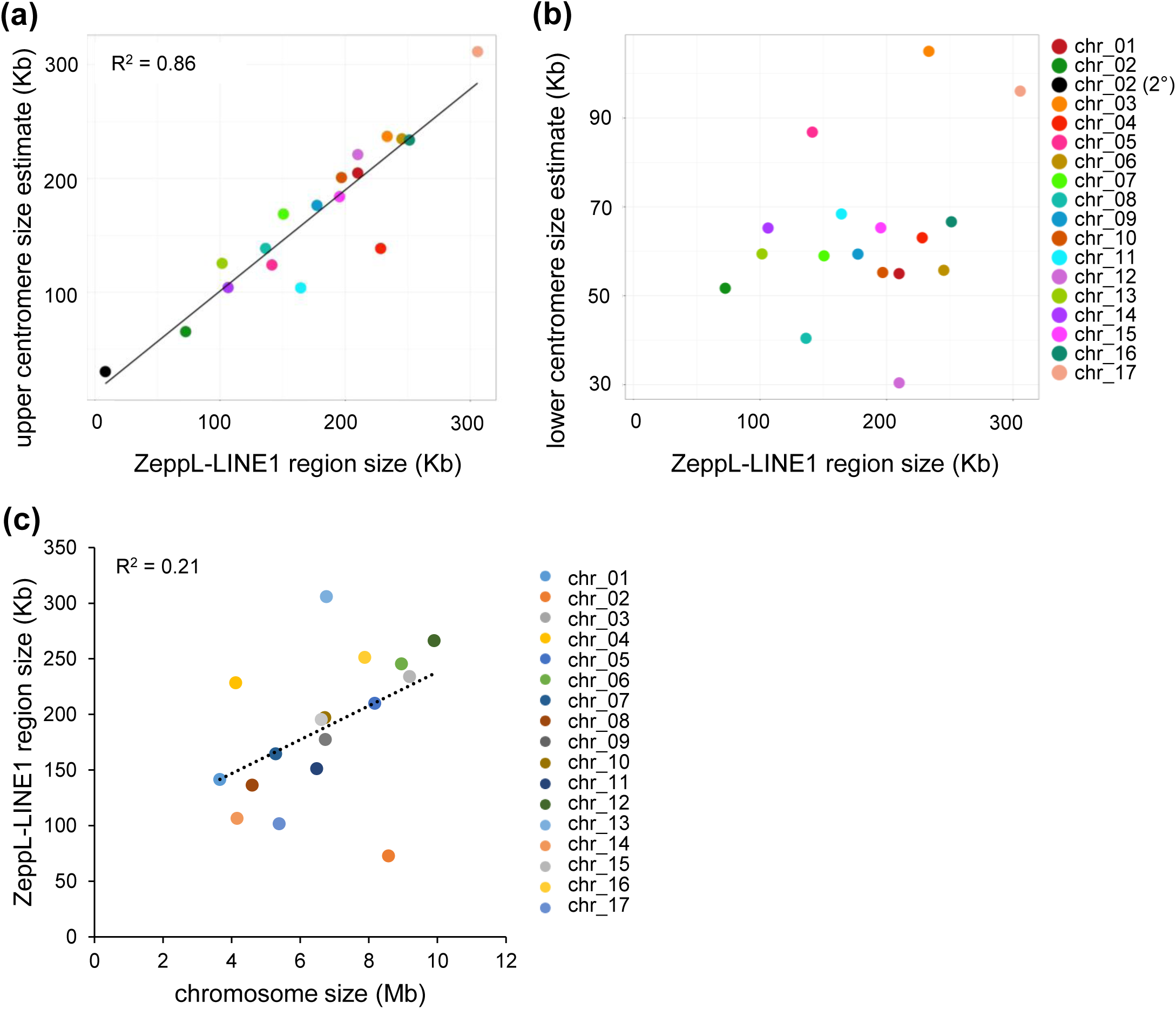
Relationships between ZeppL-enriched region size, centromere size and chromosome size for Chlamydomonas strain UL-1690. **(a)** Upper centromere size estimates plotted against the size of ZeppL enriched regions on each chromosome with linear correlation and R^2^ value. **(b)** Lower centromere size estimate plotted against ZeppL enriched region sizes shows no correlation. **(c)** ZeppL repeat region size plotted against chromosome size shows a weak positive correlation. See Materials and Methods for details on size estimations of centromeres and ZeppL repeat regions.

**Figure S6.**
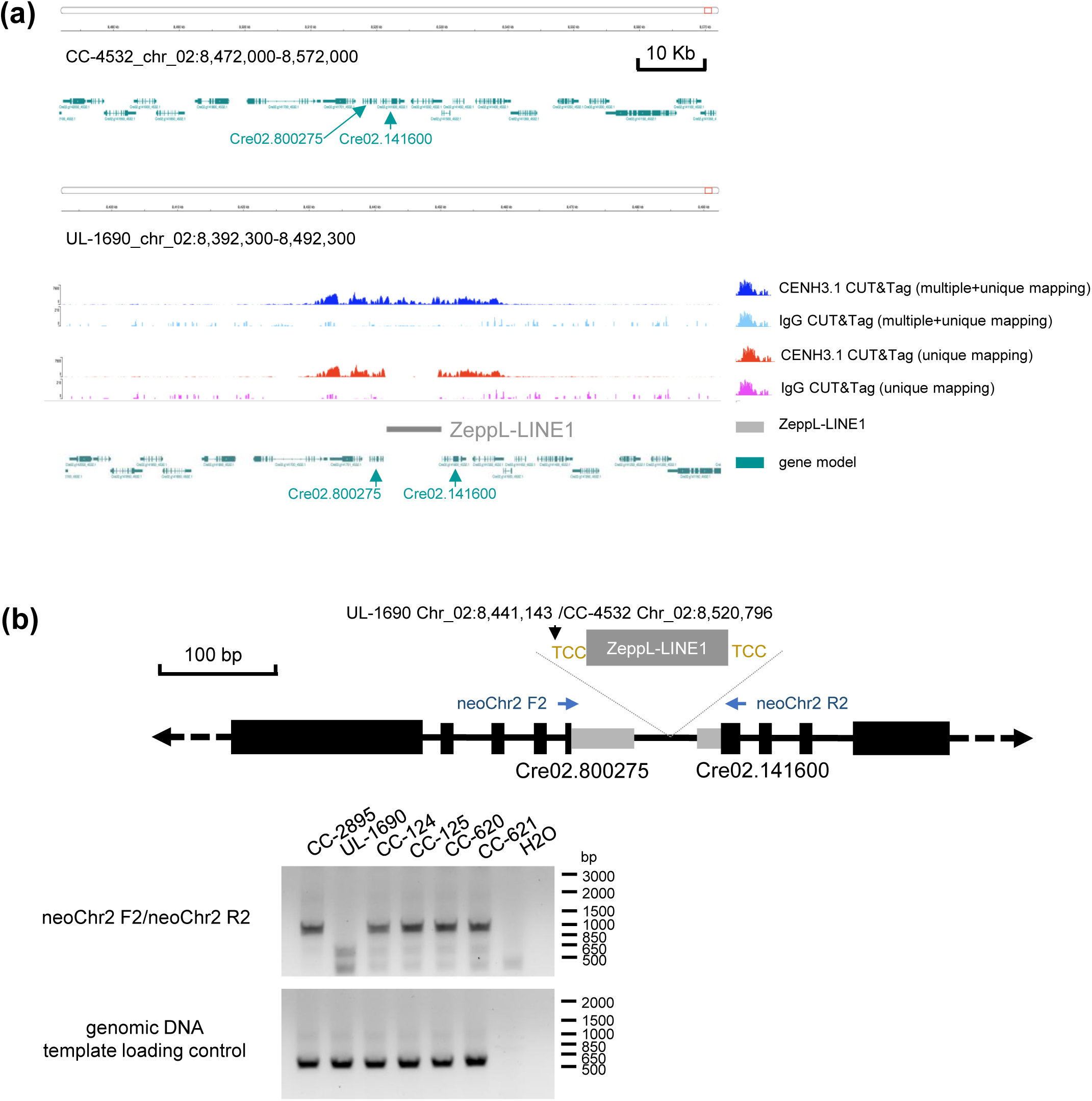
Genome browser view of CENH3 and IgG CUT&Tag read distribution around the chr2 neocentromere region. **(a)** Upper browser panel shows CC-4532 genome at neocentromere region without the ZeppL insertion. Lower panel shows the UL-1690 genome at the neocentromere site with the ZeppL insertion and mapping data for CUT&Tag reads. Annotation is the same as Figure 4. **(b)** Genotyping the junction region on chr2 flanking the ZeppL insertion on chr2. Upper panel shows a schematic of insertion between genes Cre02.g800275 and Cre02.g141600 with genotyping primer locations indicated. ZeppL fragment (8334bp) schematic is not scaled to actual size. Upper gel shows PCR products using the genotyping primers depicted in the above diagram. Lower panel shows genotyping of the same samples with primers for the *CENH3.1* locus (CENH3.1 F1/CENH3.1 R1 in Table S2) as a template quality control.

**Figures S7.**
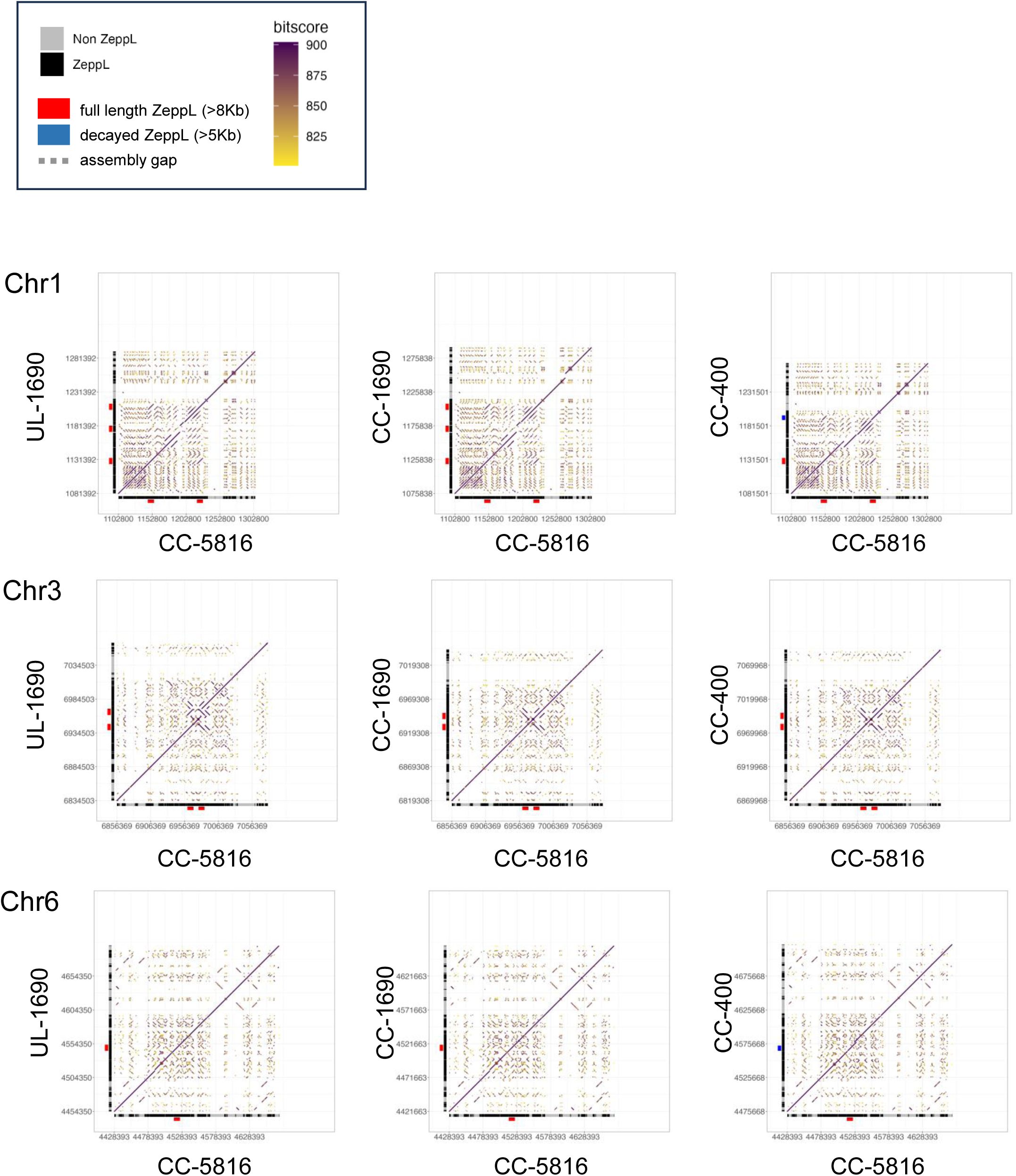
Dot plot comparisons of centromere regions of chromosomes containing one or more full length (>8kb, red bars) or partially decayed (>5kb, blue bars) ZeppL elements analyzed in Figure 5. Black blocks and grey blocks along each axis represent ZeppL or non-ZeppL sequences, respectively. Centromere coordinates are listed in Table S6.

**Figures S8.**
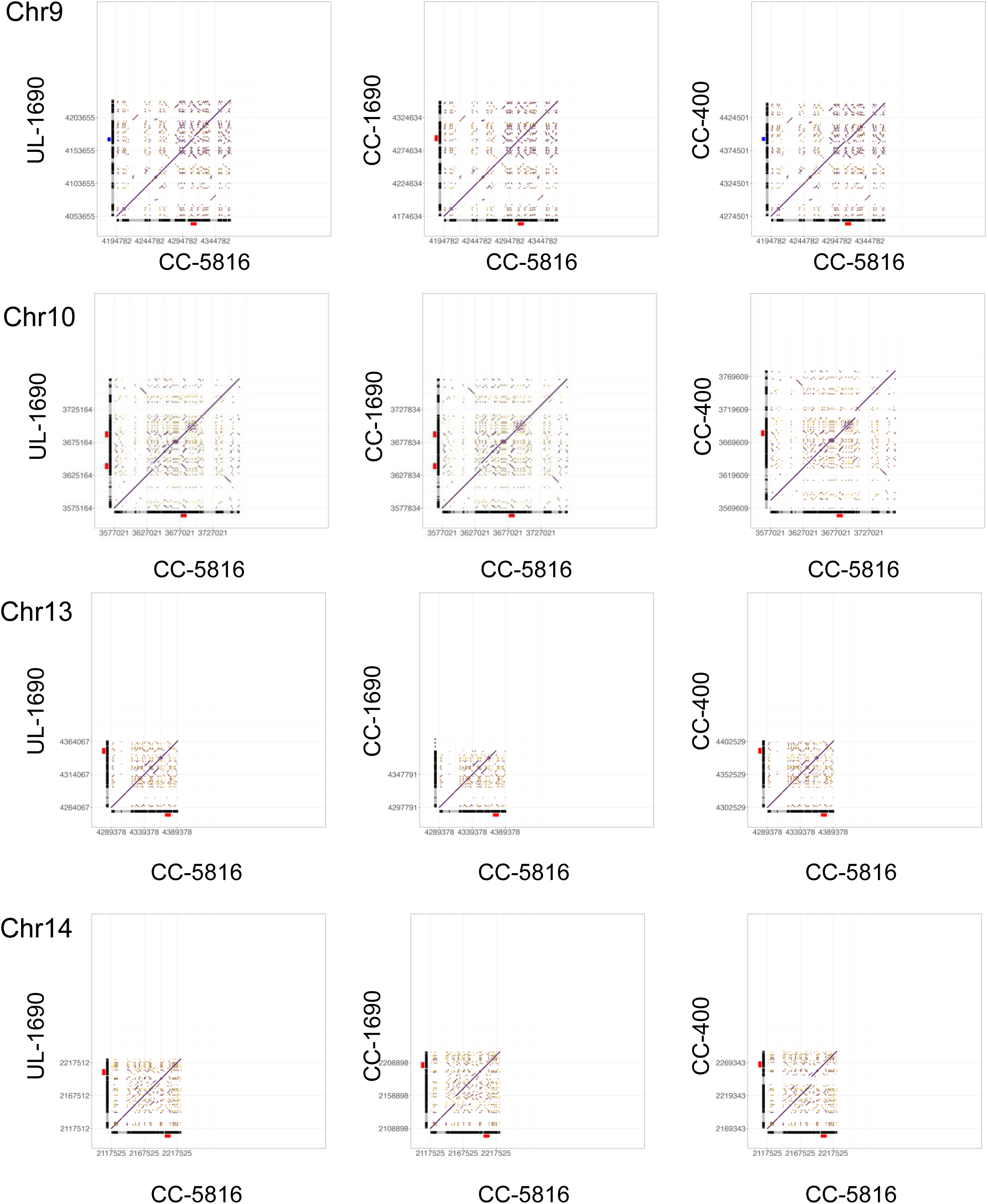
Dot plot comparisons of centromere regions of chromosomes containing one or more full length (>8kb, red bars) or partially decayed (>5kb, blue bars) ZeppL elements analyzed in Figure 5. Black blocks and grey blocks along each axis represent ZeppL or non-ZeppL sequences, respectively. Centromere coordinates are listed in Table S6.

**Figures S9.**
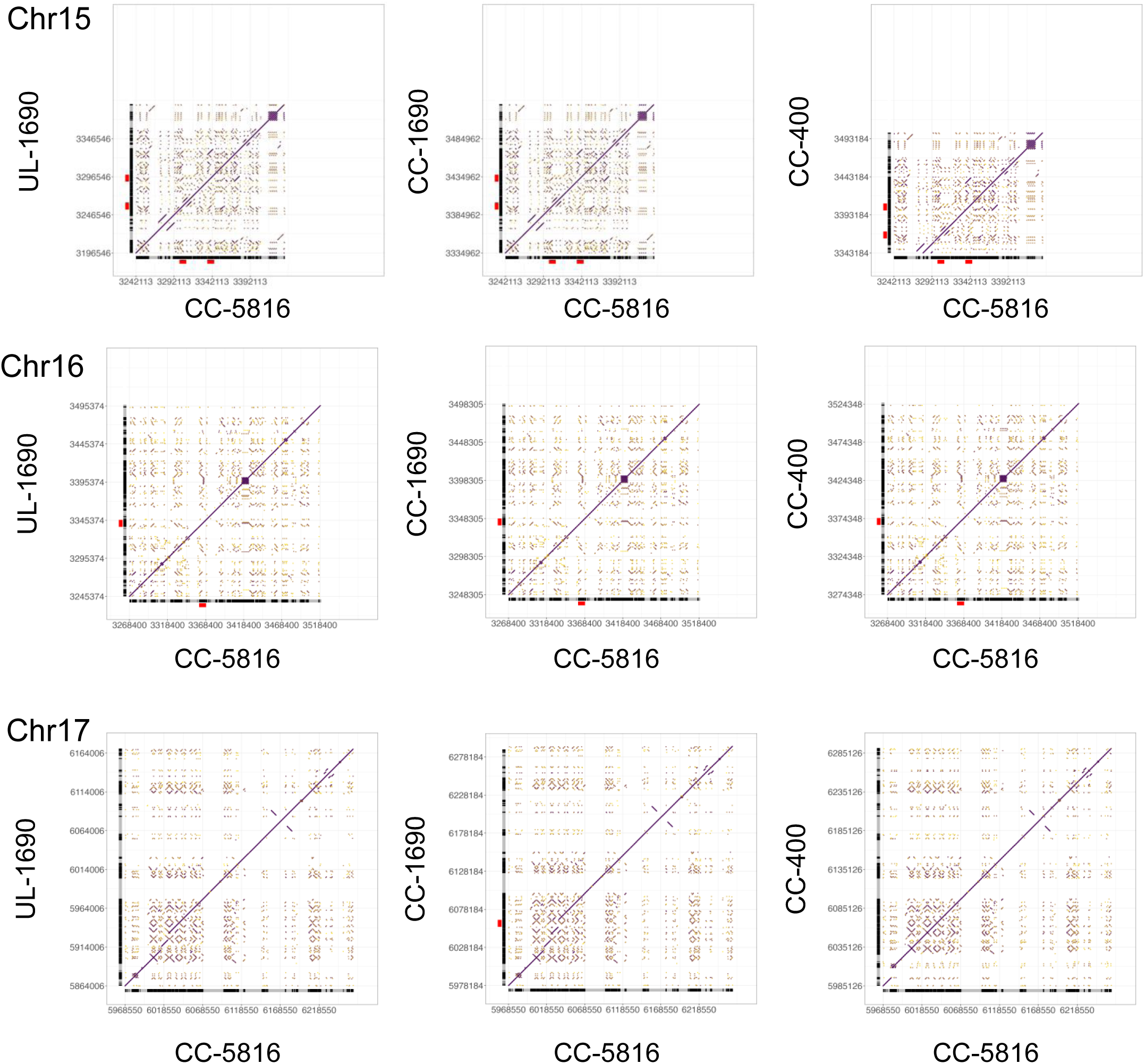
Dot plot comparisons of centromere regions of chromosomes containing one or more full length (>8kb, red bars) or partially decayed (>5kb, blue bars) ZeppL elements analyzed in Figure 5. Black blocks and grey blocks along each axis represent ZeppL or non-ZeppL sequences, respectively. Centromere coordinates are listed in Table S6.

**Figures S10.**
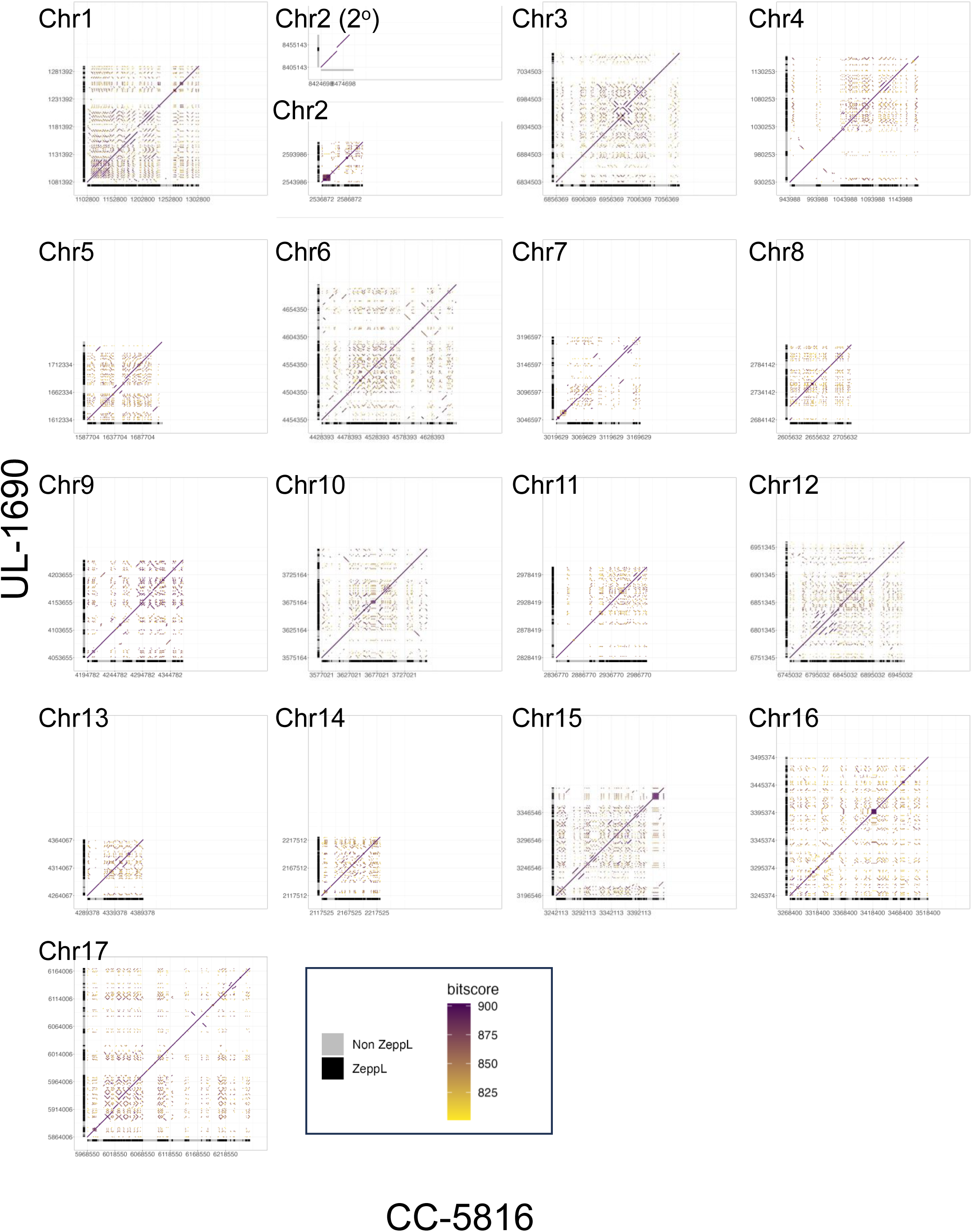
Dot plot comparisons of all CC-5816 centromeres (x axes) against corresponding centromere regions on y axes from UL-1690; this study (Figure S9), CC-1690; (O’Donnell et al., 2020) (Figure S10) and CC-400; this study (Figure S11). CC-5816 data are from (Payne et al. 2023). These plots provide a comprehensive comparison of all centromere regions.

**Figures S11.**
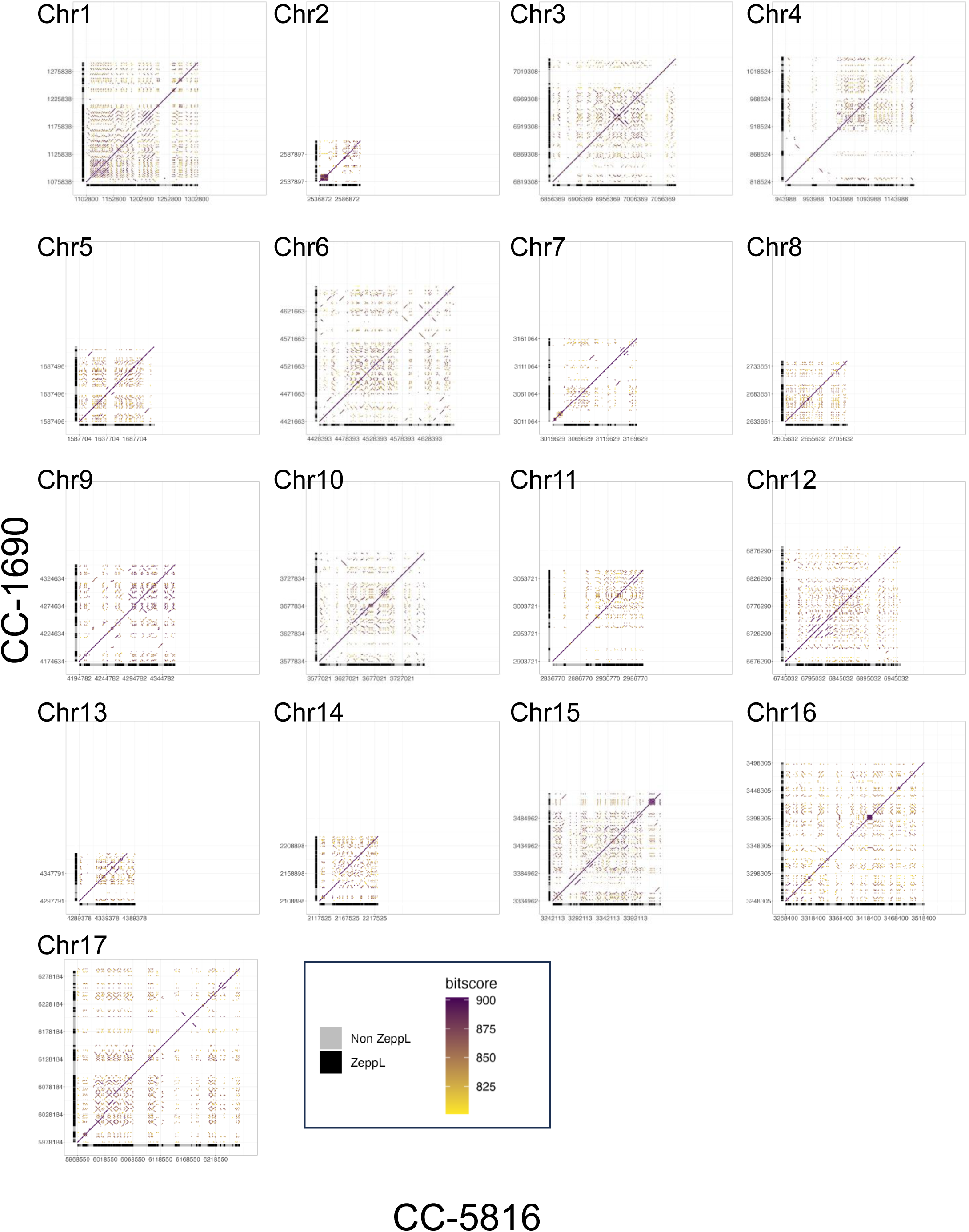
Dot plot comparisons of all CC-5816 centromeres (x axes) against corresponding centromere regions on y axes from UL-1690; this study (Figure S9), CC-1690; (O’Donnell et al., 2020) (Figure S10) and CC-400; this study (Figure S11). CC-5816 data are from (Payne et al. 2023). These plots provide a comprehensive comparison of all centromere regions.

**Figures S12.**
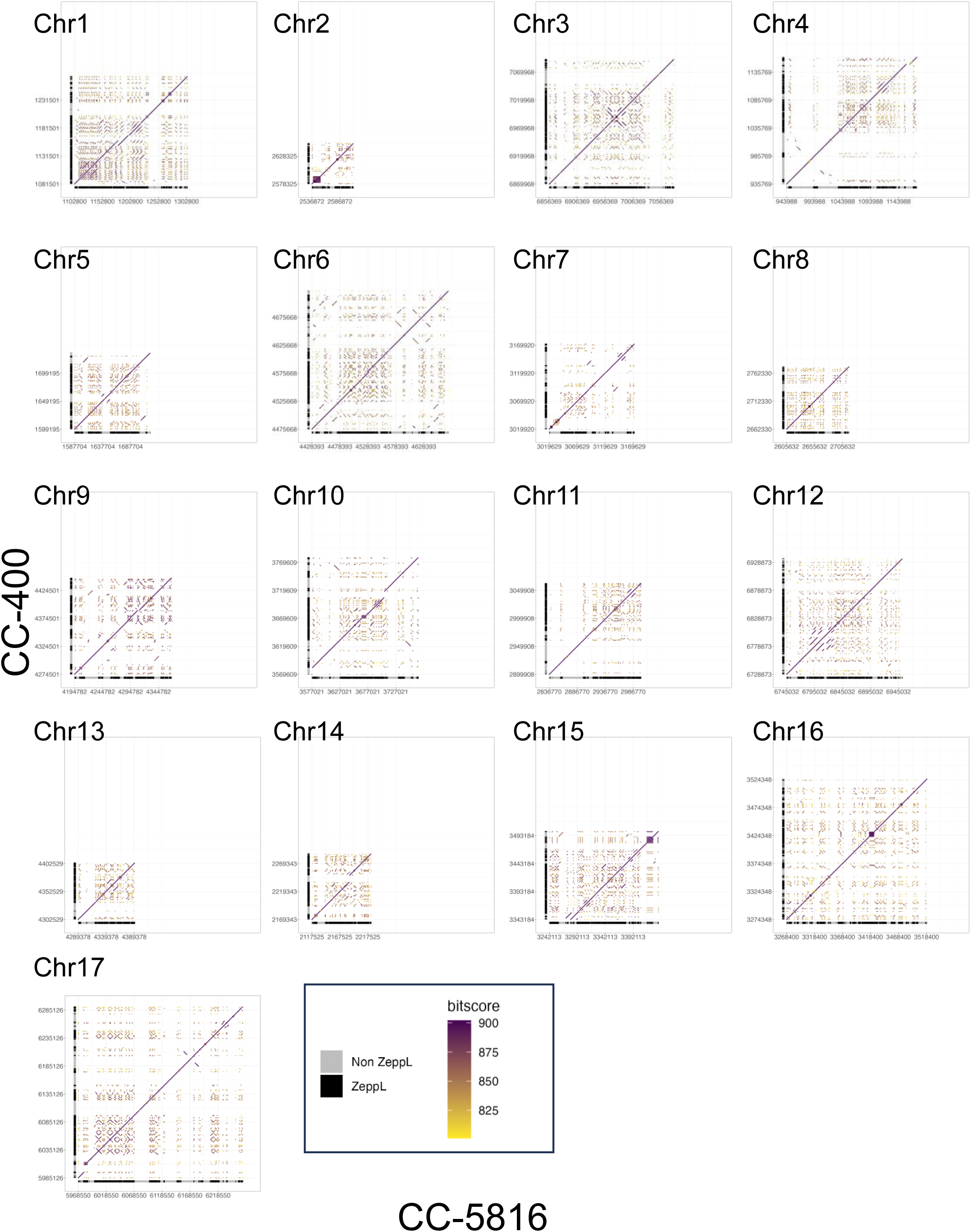
Dot plot comparisons of all CC-5816 centromeres (x axes) against corresponding centromere regions on y axes from UL-1690; this study (Figure S9), CC-1690; (O’Donnell et al., 2020) (Figure S10) and CC-400; this study (Figure S11). CC-5816 data are from (Payne et al. 2023). These plots provide a comprehensive comparison of all centromere regions.

**Figure S13.**
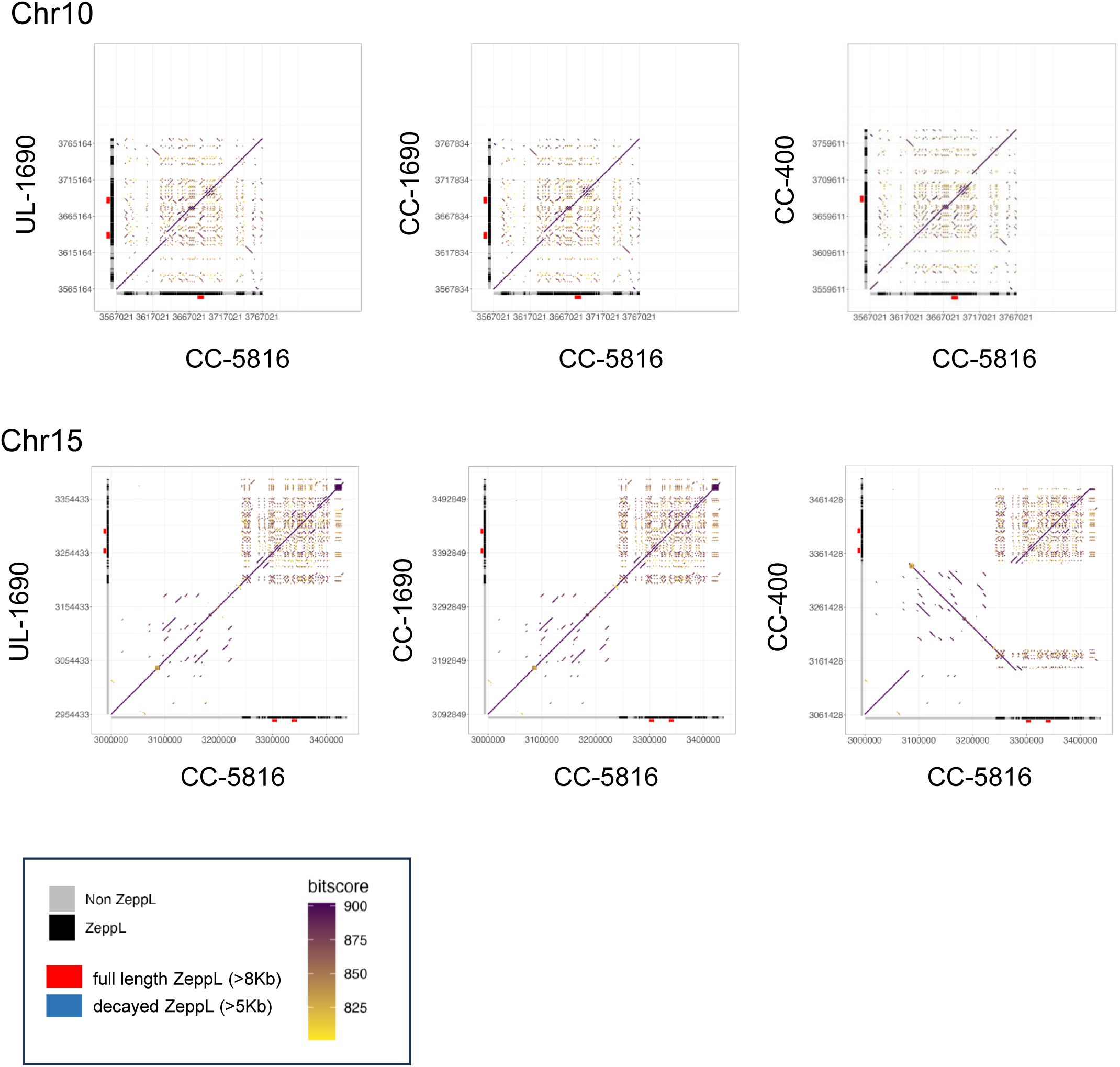
Side by side dot plot comparisons of centromere regions in chr10 and chr15 to illustrate possible structural differences in the CC-400 assembly compared to other assemblies. Annotation is the same as in Figures S10-S12, where both chr10 and chr15 of CC-400 have noticeably smaller ZeppL enriched regions. Alignment regions were extended to 200 kb on chromosome 10 and 430 kb on chromosome 15 to capture structural variations in CC-400. This extended alignment identified a ∼400kb inversion near the centromeric region of chromosome 15 in CC-400 which was confirmed manually by inspecting PacBio HiFi reads spanning the inversion junctions.

### Supporting Tables

Table S1. CRISPR-Cas9-mediated mutagenesis target site sequences and mutant junction sequences.

Table S2. Oligonucleotides used in this study.

Table S3. Gap positions in four Chlamydomonas genome assemblies.

Table S4. Tandem repeats regions in chr11 and chr15 in genome assemblies CC-5816, CC-400, and UL-1690.

Table S5. Coordinates of centromere locations and ZeppL-LINE1 regions in UL-1690 HiFi genome.

Table S6. Coordinates of ZeppL-LINE1 enriched region in UL-1690, CC-400, CC-5816, and CC-4532 assemblies.

Table S7. Coordinates and percentage identity of partially decayed ZeppL-LINE1 elements to the corresponding full length ZeppL elements in CC-5816.

### Supporting Data

**Data S1.** Sequences of ZeppL-LINE1 enriched regions in four genome assemblies.

**Data S2.** Sequences of full-length ZeppL-LINE1 elements (>8kb) in four genome assemblies.

**Data S3.** Coordinates and percent identity of partially decayed ZeppL-LINE1 elements to the corresponding full length ZeppL fragments in CC-5816 shown in Figures 5, S7.

## Notes

https://github.com/dawelab/Chlamydomonas_centromeres

### Competing Interest Statement

The authors have declared no competing interest.

